# Basal ganglia-independent thalamic bursts do not wake cortex during sleep

**DOI:** 10.1101/2025.08.10.669514

**Authors:** Xiaowei Liu, Jing Guang, Zvi Israel, Denise Wajnsztajn, Aeyal Raz, Hagai Bergman

**Author notes:** These authors contributed equally to this work (Xiaowei Liu, Jing Guang). These authors are corresponding authors.

## Abstract

The thalamus is a key forebrain structure that gates peripheral, subcortical, and cortico-cortical communication^1,2^. Awake thalamic bursts provide the cortex with a “wake-up” signal^2–4^. Paradoxically, thalamic neurons discharge tonically during cellular depolarization and activated brain states (wakefulness, REM sleep) but burst during hyperpolarization and NREM sleep^5–9^. It has been proposed that NREM thalamic bursts do not awaken the cortex because of their periodic and synchronized nature^2–4^; however, this has never been tested *in vivo* during natural sleep. We simultaneously recorded polysomnographic signals, local field potentials, and spiking activity from multiple thalamic neurons in the ventral anterior and centromedian nuclei of two female non-human primates during naturally occurring vigilance states. These nuclei receive GABAergic output from the basal ganglia^10,11^, with discharge rate and GABA outflow decreasing during NREM sleep^12^. We found that despite the expected thalamic depolarization, bursting increased significantly. NREM bursts were neither periodic nor highly synchronized. However, EEG activity time-locked to burst onset during NREM sleep differed markedly from that observed during wakefulness and REM sleep. These results support a modulatory, rather than a driving, relationship between the basal ganglia and thalamus. NREM thalamic bursts do not awaken the cortex, probably due to unique state-dependent thalamocortical dynamics.

---

The classical D1/D2 direct/indirect model and other computational models of the basal ganglia (BG)^10,11^ assume that the BG output structures, the internal segment of the globus pallidus (GPi) and the substantia nigra pars reticulata (SNr), drive their thalamic targets, the ventral anterior (VA), ventral lateral (VL), centromedian (CM), and parafascicular (Pf) nuclei. The thalamic neurons relay the BG inhibitory or disinhibitory drive to the frontal cortex^10,11^, as well as to the BG input structures (e.g., striatum)^13^. Uniformly, BG physiologists assume a driving relationship between the BG and their thalamic targets. On the other hand, thalamic scholars often categorize thalamic nuclei into two types^1,2,14,15^, e.g., first-order nuclei, which are driven by peripheral or subcortical inputs, and higher-order nuclei, which are driven by cortical inputs from layer 5^2^. They point to the GABAergic nature of the BG-thalamus connectivity and treat the VA, VL, and CM as higher-order thalamic nuclei modulated by their BG input^2,14,16^.

In addition to the effects of peripheral and subcortical inputs, the activity of both the thalamus and cortex is significantly modulated by brainstem and basal forebrain cholinergic and monoaminergic neuromodulation across the vigilance states — wakefulness, rapid eye movement sleep (REM), and non-rapid eye movement sleep (NREM)^17^. Schematically, there is a high tone of all neuromodulators during wakefulness and a reduction during NREM sleep. REM sleep differs, with a further decrease in monoaminergic drive but a resumption of cholinergic tone to a similar level as in the awake state^18^.

*In vivo* studies have demonstrated that thalamic neurons fire in tonic mode during wakefulness and REM sleep, but switch to burst mode during NREM sleep^2,5–7^. The high density of low-threshold calcium channels in the thalamocortical cells, and the changes in the resting membrane potentials of these neurons, are the root cause of this behavior^8,9^. During NREM sleep, the glutamatergic drive (e.g., from the retina to the lateral geniculate nucleus) of thalamocortical neurons decreases, leading to hyperpolarization and a high tendency to burst. Alternatively, it may be a reduction in the cholinergic modulation that results in NREM thalamic hyperpolarization and bursting^19,20^. The situation is reversed in the awake state; the thalamocortical neurons are depolarized, fire tonically, and reliably encode the peripheral inputs^2,3^.

However, thalamic bursts were also observed in the awake state^3,4^. The post-synaptic effects of bursts are more influential than those of single spikes^21,22^. Therefore, bursts can be used to decode salient events and “wake-up calls” for the cortex^4^. This naturally raises the question of why the NREM thalamic bursts do not wake the cortex. The textbook answer is that the rhythmic and synchronized bursts of thalamocortical cells during NREM sleep reflect a different message for the cortex, which indicates a complete lack of information^2^. This hypothesis is inspired by *in vitro* demonstration of periodic and synchronized bursts of hyperpolarized neurons^8,9^. However, the periodicity and synchronization of thalamic bursts have never been analytically examined (for periodicity) or experimentally tested (for synchronization) in *in vivo* conditions.

Here, we recorded the VA and CM electrophysiological activity, in parallel with polysomnography, across the three vigilance states of two non-human primates. These nuclei are innervated by the GPi and SNr, whose discharge rate is reduced during low arousal and NREM sleep states^12,23^. This is likely associated with a reduced GABA release, which may lead to depolarization of VA and CM neurons during NREM and hyperpolarization during wakefulness and REM sleep. This would reverse the typical relationship between vigilance states and thalamic firing modes: tonic firing would occur during NREM sleep, and bursting would emerge during wakefulness and REM sleep. This pattern is opposite to that observed in thalamic nuclei driven by glutamatergic input, such as the lateral geniculate nucleus, where tonic firing predominates during both wakefulness and REM sleep, and bursting is characteristic of NREM sleep. Therefore, the primary objectives of this study were to elucidate the tonic-burst activity of the BG-related thalamic nuclei and to understand why NREM thalamic bursts do not awaken the cortex. The results led to a paradigm shift in our understanding of the relationships between BG, thalamus, and cortex.

## Results

Two female African green monkeys (Chlorocebus aethiops or sabaeus, weight: 4-5 kg, monkey Md and Wh) contributed to the experiments (Fig. 1a-d). All experimental procedures were conducted in accordance with the National Institutes of Health Guide for the Care and Use of Laboratory Animals and the ethical guidelines of the Hebrew University of Jerusalem. Before the experimental recording, the animals were habituated to sleeping in the lab. A head holder, bilateral eye coils, two frontal skull-mounted EEG electrodes, and a 27 × 34 mm recording chamber located above a central frontoparietal craniectomy site were implanted across four separate surgeries. Surgeries were performed under general anaesthesia with appropriate antibiotic and analgesic administration. Each surgery was followed by a recovery and re-training period of 3-6 weeks. Finally, a 3T MRI scan verified the precise location of the recording chamber (Fig. 1a).

**Figure 1.**
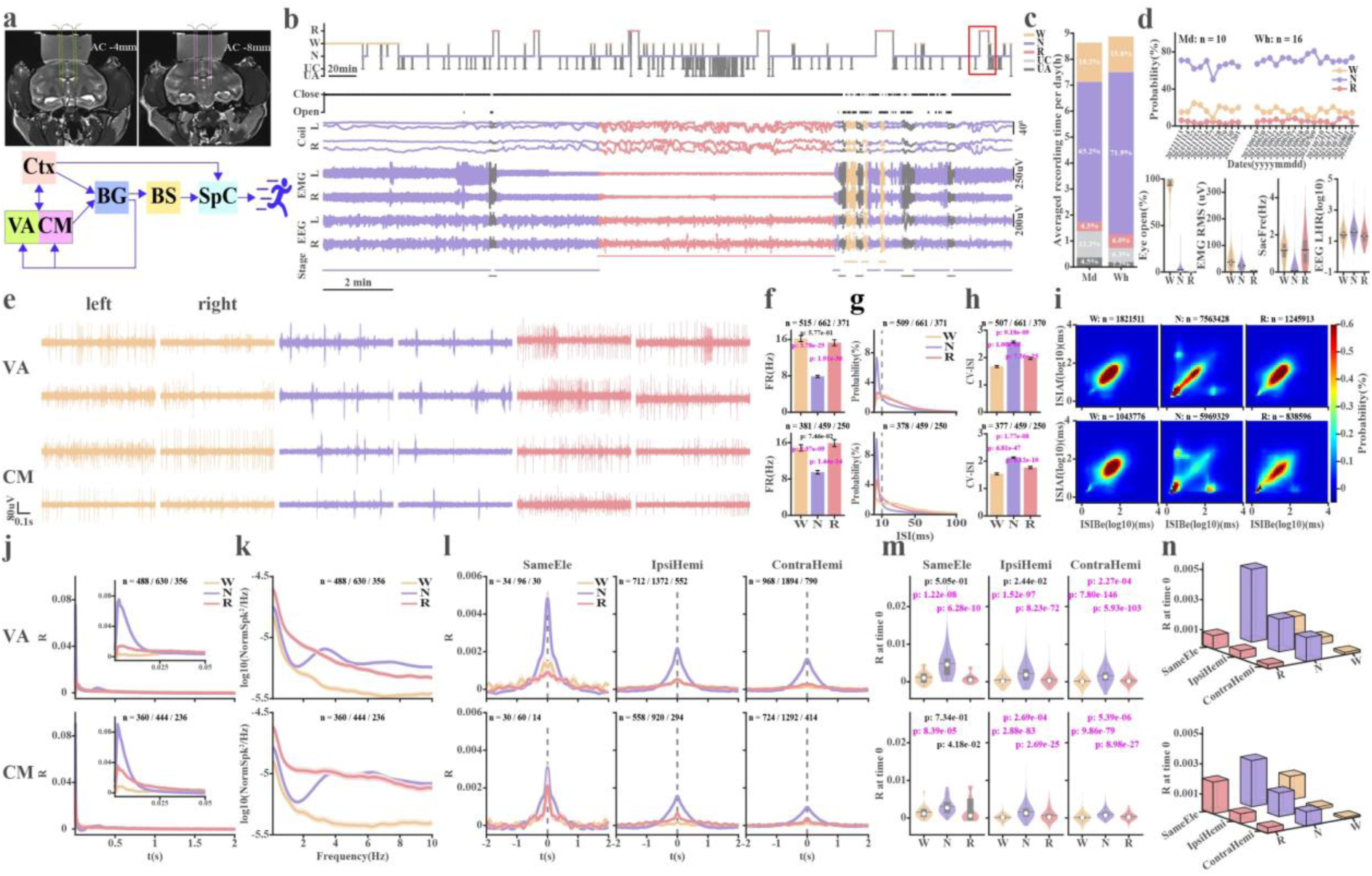
Experimental methods, sleep architecture, and general thalamic physiology across vigilance states. **a**, MRI and anatomical scheme of thalamic recording targets and their connectivity. **b**, One-night hypnogram and 20-min (marked in red square) polysomnographic examples. The awake (W), NREM (N), REM (R), and unclassified/unavailable (UC/UA) epochs are marked in yellow, purple, pink, and light and dark grey, respectively. **c**, Sleep architecture of the two monkeys: the mean duration and percentage of total time in each wake-sleep state. **d**, top: proportion of W, N, and R states per session across sequential recording nights in both NHPs (the nights displayed here only include those for which the recorded neurons are all from the thalamus). Bottom: Violin plots of polysomnographic metric distributions across vigilance states. The number of 10-second epochs for W, N, and R is 13,343, 55,350, and 4,320, respectively. **e**, Examples of W, N, and R spiking activity simultaneously recorded by two microelectrodes (upper and lower subplots) in the left and right hemispheres. **f**, Firing rate (FR) of thalamic neurons. **g**, Inter spike intervals (ISI) histograms. The dashed vertical grey lines mark ISI of 10 ms. **h**, Coefficient of variation of ISIs (CV-ISI). **i**, ISI return maps of ISI (n+1, after, ISIAf) vs ISI (n, before, ISIBe). **j**, Auto-correlation histograms of spike trains (lag = 2 s). The insets depict the auto-correlation histograms near time zero (lag = 50 ms). **k**, Power spectrum densities of spike train. **l**, Cross-correlation histograms of spike trains (lag = ±2 s). **m**, The distribution of R values at time zero. The number of time-zero R values equals the number of neuron pairs with repetition shown on the subplots in **l**. **n**, The average time-zero R values as a function of the vigilance state and the distance. Data are shown as mean ± SEM (standard error of mean) in **f**, **h**, and **j**-**l**. The n indicates the number of days in **d**, neurons in **f**-**h** and **j**-**k**, neuron pairs with repetition in **l**, and ISI pairs in **i**. A Bonferroni-corrected Wilcoxon rank sum test was used to calculate the statistically significant difference in FR, CV-ISI, and R values at time 0 among different sleep stages (p < 0.05/3). The p-values indicating significant differences are marked in magenta; otherwise, they are in black. Abbreviations. BG-basal ganglia; BS-brain stem; CM-centromedian nucleus; Ctx-cortex; ContraHemi-contralateral hemisphere; EMG-electromyography; EEG-electroencephalogram; IpsiHemi-ipsilateral hemisphere; N, NREM-Non-rapid eye movement; R, REM-Rapid eye movement; RMS-root mean square; SameEle-same electrode; SpC-spinal cord; VA-Ventral anterior thalamic nucleus. Colour coding: Yellow-Wake, Purple-NREM, Pink-REM.

Recordings were conducted in a noise-attenuated room. The experiment was usually started at 4 PM. The light in the room was turned off at around 7-8 PM, and recording usually continued until 4-5 AM when the monkey returned to the open yard in the animal facility with her peers. The eye-open/close state and polysomnography were monitored (Fig. 1b-d). Additionally, we recorded local field potential (LFP) and spiking activity from two sets of four microelectrodes targeting the right and left VA or CM thalamic nuclei.

Here, we report the results of VA and CM neuronal activity (1161 neurons with isolation quality^24^ > 0.7 for at least 60 seconds), recorded over 37 nights in two NHPs. Table S1 provides a detailed description of the neuronal database. The results of the subclass of neurons with better isolation scores (>0.85, Table S1) are reported in Fig. S1 and align with those reported here. The results of the two NHPs did not differ (Fig. S2) and were therefore pooled for all subsequent analyses.

### Sleep architecture

Figure 1b illustrates a typical hypnogram and a 20-minute segment showing the changes in eye movements, EMG, and EEG across vigilance states. These states were semiautomatically detected^12^ for non-overlapping 10-second periods. A small fraction of the data was not classified, either due to data loss or ambiguity, and was therefore discarded from the analysis. Figure 1c shows the average fraction of the vigilance states. The sleep patterns and their characteristics remained stable throughout the experiment duration (Fig. 1d).

### Spiking activity in the thalamus across vigilance states

VA and CM neurons depict the expected tonic/burst/tonic activity during the awake, NREM, and REM sleep states, respectively (Fig. 1e). The average discharge rate of VA and CM neurons was approximately 15 spikes/s in the awake and REM sleep states (Fig. 1f). In line with previous thalamus studies^5–7,25^ and other brain structures^12^, the firing rate during NREM sleep significantly decreased (Fig. 1f).

We used several methods to quantify the firing pattern of VA and CM neurons. Fig. 1g shows that the inter-spike interval (ISI) histograms during wakefulness and REM are relatively flat, while short ISIs (<10 ms, characteristic of burst firing) during NREM are markedly more frequent. The coefficient of variation of the ISIs (CV-ISI) for the thalamic neurons range from 1.5 to 2.5 (Fig. 1h). CV-ISI values are significantly higher during NREM compared to wakefulness and REM, with REM showing slightly elevated CV-ISI values relative to wakefulness (Fig. 1h and Fig. S1c). The ISI return maps (Fig. 1i and Fig. S1d) reveal bursting activity during NREM. The initial peaks of the average autocorrelation histograms are more pronounced during NREM (Fig. 1j). Finally, the average power spectrum densities (PSD) of thalamic spike trains (Fig. 1k) do not exhibit narrow peaks.

Neuronal synchronization is best measured by cross-correlation analysis. In this study, neuron pairs were simultaneously recorded by the same electrode, or by different electrodes in the ipsilateral or contralateral hemispheres. The distances between the corresponding neuron pairs in these situations are ≤ 0.2^26^, 0.5-4, and 5-10 mm, respectively. The average correlation coefficient (R) cross-correlation histograms of all spikes (Fig. 1l-n) depict a symmetric positive correlation around time 0 in NREM, suggesting a “common-input” mechanism^27^. Fig. S3 shows that the conditional discharge rate correlation histograms reveal similar properties. We also calculated the correlation histograms of the multi-unit activity (i.e., raw signals filtered by a 300 - 9000Hz bandpass). The results (Fig. S4) align with the above reports.

### Detection of thalamic bursts and their properties across vigilance states

Our raw data (Fig. 1e) and discharge pattern analysis (Figs. 1g-k, S1b-d, i, j, S3a, and S4a) suggest that VA and CM thalamic neurons tend to burst more frequently during NREM. We, therefore, tested several burst detection methods^28^ to identify bursts and to investigate their properties (Fig. 2). We found that the maximal-interval method best matched the expert choice and used it hereafter. The Poisson surprise method^29^ yielded more conservative, but similar results (Fig. S5).

**Figure 2.**
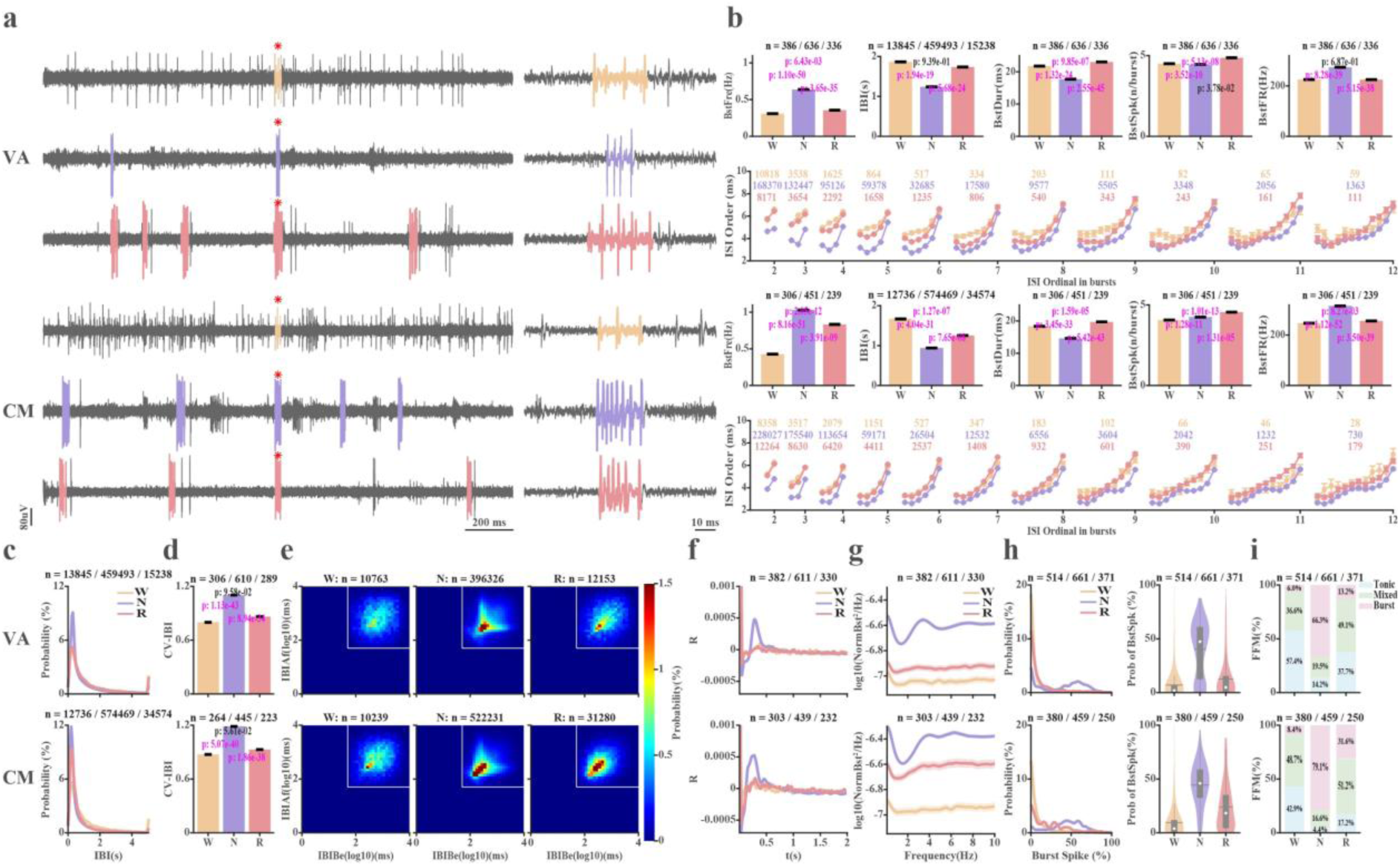
Burst detection, features, and pattern across the wake-sleep stages. **a**, Examples of burst detection. In each thalamic nucleus, 2-second traces are shown on the left and 100 ms segments around the burst marked by a red star are displayed on the right (detected bursts are coloured). **b**, burst features for the VA and CM (two upper and two lower rows, respectively). Row 1^st^, from left to right: Burst frequency (BstFre), Inter-burst interval (IBI), burst duration (BstDur), number of spikes per burst (BstSpk), intra-burst discharge rate (BstFR). Row 2^nd^ ordinal inter-spike intervals in bursts of different lengths. Only bursts with fewer than 12 intervals are shown. **c**, IBI-Time Histograms. **d**, Coefficient of variations of the IBIs (CV-IBI). **e**, IBI return map (IBIAf, IBI (n+1) vs. IBIBe, IBI(n)). White lines indicate log10(50) ms (the minimum silent time before a burst in the maximum interval burst detection methods). **f**, Average autocorrelation histograms of thalamic bursts (lag = 2 s). **g**, Power spectral density (PSD) of thalamic bursts. **h**, Left-Histograms of burst spikes (BstSpk) fraction in 10-second segments. Right - the distribution of burst spikes fraction per 10-second spike trains. **I**, fraction of firing modes (FFM, burst, tonic, and mixed) in different vigilance states. Data are shown as average ± SEM in **b**, **d**, **f**, and **g**. The n indicates the number of IBIs in subplot **b**-2^nd^ column and **c**, IBI pairs in **e**, bursts in 2^nd^ and 4^th^ rows of **b** and neurons in the rest subplots. A Bonferroni-corrected Wilcoxon rank sum test was used to calculate the statistically significant difference of burst features among different vigilance stages (p < 0.05/3) in **b** and **d**. The p-values indicating significant differences are marked in magenta; otherwise, they are in black. Abbreviations and color coding as in Fig. 1.

Figs. 2a and S5a illustrate the raw spiking data with detected bursts marked in color. The burst properties are shown in Figs. 2b, S1e and S5b. The mean burst frequency ranges from 0.4 to 1 bursts/s, and is higher during NREM than during wakefulness and REM sleep. Accordingly, the mean inter-burst interval (IBI) varies between 1 and 2 seconds and is shorter during NREM sleep. This long IBI is in line with the low-threshold calcium mechanism and previous findings^5–7^. The mean duration of bursts is 15-20 ms. It is slightly longer in the VA relative to the CM, and tends to be shorter during NREM compared with wakefulness and REM sleep. The mean number of spikes per burst is between 4 and 5 spikes. Accordingly, the average frequency of the spikes in a burst is above 200 spikes/s, which is more than 10 times the tonic firing rate. Fig. S6 details the tonic activity (after burst removal) for comparison with the thalamic overall spiking activity (Fig. 1), and the burst activity (Figs. 2 and 3). Finally, the ordinal ISI relations reveal that the bursts are decelerating (Fig. 2b, 2nd and 4th rows), in line with previous *in vivo* reports^30^ and the associations of thalamic bursts with low-threshold calcium channels and spikes^8,9^.

**Figure 3.**
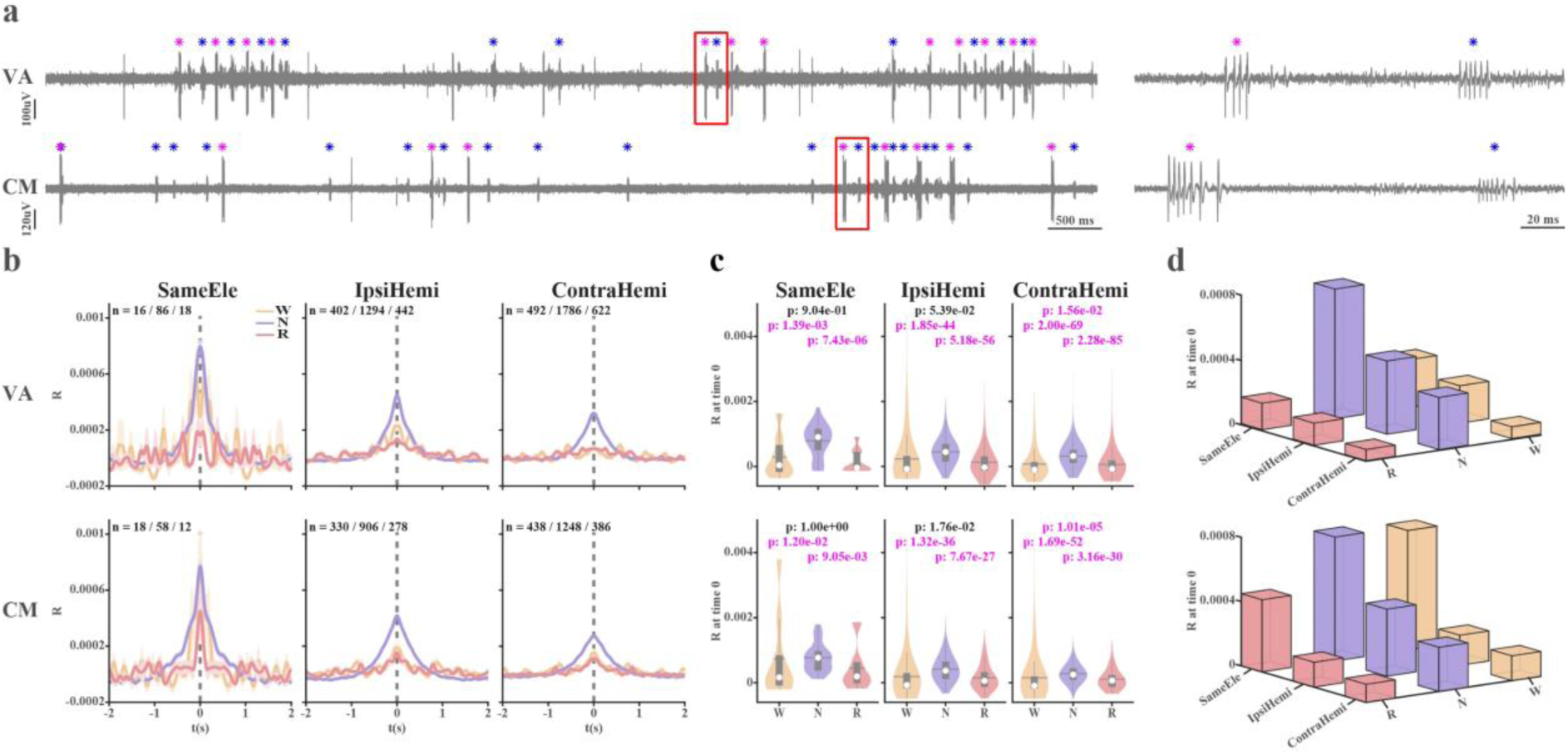
Thalamic bursts during sleep exhibit prolonged timescales and loose synchronization. **a**, An example of bursts of two units (detected bursts are marked in blue and magenta stars, respectively) simultaneously recorded by the same electrode in the VA and CM nuclei. Left – 10-second trace, Right - higher magnification of the red box area (200-ms trace). **b**, Cross-correlation histograms of bursts from the same electrode, or different electrodes in the ipsilateral or contralateral hemisphere (left to right). **c**, The distribution of R values at time 0. The number of R values at time 0 is the same as that of the neuron pairs with repetition shown in **b.** Data is shown in mean ± SEM and the n indicates the number of neuron pairs with repetition in **b**. The Wilcoxon rank sum test was used to calculate the statistically significant difference of R values at time 0 among different vigilance stages (p < 0.05/3). P values indicating significant differences are marked in magenta. Otherwise, they are in black. **d**, 3D-bar plots of the average time-zero R values as a function of wake-sleep state and distance. Abbreviations and color coding as in Fig. 1.

To furtherly explore the characteristics of thalamic bursts, we proceeded to test their temporal patterns. We marked each detected burst as a single event. These marked events, collectively referred to as the burst train, were then analyzed using the same methods previously applied to individual spikes. The IBI histograms exhibit a broad peak near zero interval duration, indicating a predominance of relatively short IBI (Fig. 2c and S1f). The CV-IBI values (Figs. 2d and S1g) range between 0.8 and 1.2, suggesting that bursting timing more closely resembles a Poisson process than a periodic pattern. NREM exhibits the highest CV-IBI (>1) among the three vigilance states. Similarly, the IBI return map, the autocorrelation histogram, and the PSD of the bursts (Figs 2e-g, S1h-j, and S7a) also did not show a robust signature of periodic activity^31,32^. Rather, these metrics indicate a tendency for temporal clustering of the bursts (bursts of bursts), particularly during NREM sleep.

The tendency for clustering of bursts suggests that there is one hierarchical level higher than the single-burst discharge. This may be related to the division of thalamic activity into tonic and burst mode states. The neural activity was segmented into 10-second epochs, aligned with the corresponding segments of polysomnography. For each segment, the fraction of burst spikes relative to total spikes was calculated (Figs. 2h, i, and S1k, l). Based on this calculation, the segments were classified into three discharge modes: burst (>30% of spikes in bursts), tonic (<3%), or mixed (3-30%). Figures 2h and S1k depict the distribution of these segments in our recordings. NREM differed markedly from the awake and REM states, with more than 65% of segments in burst mode (Figs. 2i and S1l).

### Synchronization patterns of thalamic bursts across vigilance states

Although coupled periodic processes tend to synchronize, synchronization can be detected independently of periodicity^33^. We calculated the cross-correlation histograms of burst trains for neuron pairs simultaneously recorded by the same electrode (Fig. 3a), or by different electrodes in the ipsilateral or contralateral hemispheres (Fig. 1e). Our results, revealing burst of two distinct units recorded by a single microelectrode (Fig. 3a), align with the raw data displayed in previous studies^6,7^. Figures 3b-d, S1m-o, S5j-l, and S6h-j illustrate the cross-correlation histograms, organized by distance and vigilance states. These histograms are calculated with a 0.7 ms bin width and smoothed using a moving Gaussian kernel with an SD of 60 bins. Most cross-correlation histograms demonstrate a central, symmetric, and broad peak. However, the magnitude of these peaks at time zero, indicating tight synchronicity, is extremely low. The correlation coefficient (R (t = 0)) values do not exceed 0.001 (Fig. 3d).

We also calculated the burst cross-correlation histogram as the conditional discharge rate (Fig. S7b) and found similar results. The conditional firing rate cross-correlation histogram enabled us to estimate the proportion of bursts attributable to a common input mechanism relative to the total number of bursts. We dubbed this metric the association index (AI)^34^. The bursts’ AI values are also low (AI < 0.15, Fig. S7c-f), even though their values are higher than those of the all-inclusive (Fig. S3c-f) and burst-removed (tonic activity, Fig. S6k-o) spike trains. The AI values take into account all spikes in the central peak (from -0.5 to 0.5 s) of the cross-correlation histogram, thereby including both tightly and loosely synchronized burst pairs. We, therefore, recalculated the cross-correlation histograms with 1-s bins (and no smoothing). As expected, the R (t = 0) is higher, but not exceeding 0.4 (R^2^ < 0.16). Thus, both R and AI cross-correlation analyses indicate that thalamic bursts synchronize only loosely and over extended timescales (Fig. 3b-d, Fig. S7b-f, and Fig. S8).

### The relationship between thalamic bursts and vigilance states

Up to this point, our findings suggest that thalamic bursts during sleep are neither periodic nor robustly synchronized. We, therefore, first tested whether sleep bursts triggered wakefulness or promoted transitions to micro-arousal states^35,36^.

The burst features of every 10-second segment were aligned to the vigilance state transition (e.g., from NREM to REM sleep, Fig. S9). We found that, for most vigilance state transitions, the burst frequency before and around the transition did not significantly exceed the expected values (Figs. 4a and S9a). Changes in the other two burst features qualitatively resemble those of burst frequency, except that the actual spike number per burst is significantly more than the predicted value before the transition from NREM to REM sleep (S9b, c). Additionally, bursts don’t help maintain wakefulness (Figs. 4b-Left and S9d), but rather significantly increase the probability of NREM during sleep (Figs. 4b-middle and S9e). Finally, they don’t play a role in the change of vigilance states during REM sleep (Figs. 4b-right and S9f). In a nutshell, bursts are not related to waking up our monkeys.

**Figure 4.**
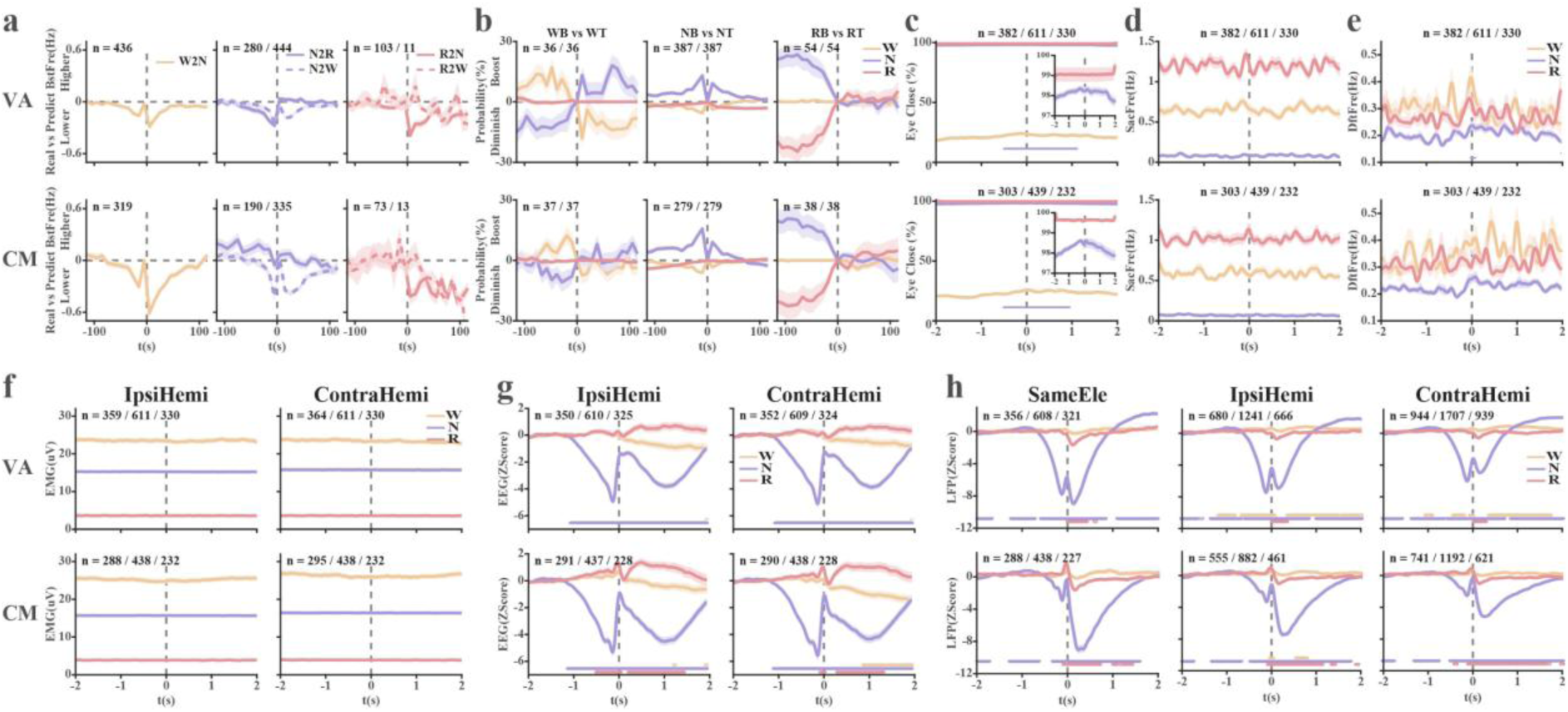
The effect of bursts on sleep stages, polysomnographic features, EEG, and thalamic LFP across awake-sleep stages. **a**, The difference between the actual and predicted burst frequencies aligned to the wake-sleep state transition. **b**, Comparison of the probability to stay in the same, or to switch to other vigilance states between the segments at time 0 with burst and tonic discharge. Positive values indicate this vigilance state is boosted or reinforced by bursts, and vice versa for negative values. **c**, Burst-triggered averages of eye open/close state. To illustrate the details of the significant difference from the baseline in NREM sleep, a zoomed-in subplot is displayed as an inset. **d-h**, Burst-triggered fast (saccadic) eye movement, slow (drift) eye movements, EMG, EEG, and LFP. The data used for the statistical analysis in **c**-**h** and the Z-Score normalization in **g** and **h** is the baseline data collected at the two-to-one-second segment before the burst event (-2 s to -1 s). All results are shown as mean ± SEM. The n means the number of neurons in **a**-**h**. The colored circles in the subplots **c**-**h** indicate statistically significant difference from the baseline (Wilcoxon rank sum test, corrected by the Bonferroni method, p<0.05/4001 in **c**-**e**, p < 0.05/5501 in **f**-**h**; sampling rate 1000 and 1375 Hz, respectively). Abbreviations. W2N, N2R, N2W, R2N, and R2W - transitions from W to N, from N to R, from N to W, from R to N, and from R to W, respectively. WB vs WT – a neuron firing in burst or tonic mode in the W state. NB vs NT – a neuron firing in burst or tonic mode in the N state. RB vs RT – a neuron firing in burst or tonic mode in the R state. The remaining abbreviations and color coding are as shown in Fig. 1.

The 10-second temporal resolution of our behavioral analysis might mask fast changes. Therefore, we examined the effect of a single burst on polysomnographic metrics with a smaller timescale (± 2 s) and high time resolution (0.7 or 1 ms). The percentage of eye closure increases around the bursts during NREM sleep (Fig. 4c). Additionally, the frequency of fast (saccade) and slow (drift) eye movements (Fig. 4d, e) as well as the EMG power (Fig. 4f) were not affected by thalamic bursts.

### The relationship between thalamic bursts and EEG/LFP activity across vigilance stages

To find hints for resolving the thalamus-sleep physiology riddle, we further examined the effects of thalamic bursts on the simultaneously recorded EEG and LFP. The EEG activity time-locked to burst onset is shown in Figure 4g. It differs across vigilance stages. During NREM sleep, EEG activity begins to decline (indicating depolarization) one second before the burst. It is followed by a rapid rebound at the burst time and subsequent negative waves. The REM EEG shows a small positive deflection before the thalamic bursts, which is more evident in CM. In the awake state, the burst triggers a small, slow positive EEG deflection, followed by a moderate, prolonged decrease after the burst. There are small differences between the awake and REM EEG responses. However, the preceding robust NREM negative EEG deflection is unmatched.

Cortical LFP is often referred to as local EEG. Monopolar subcortical LFP is probably affected by volume conductance from the cortex^37,38^, and correlates with a non-linear summation of cortical activity. Figure 4h shows the burst-triggered LFP recorded by the same electrode, or by different electrodes in the ipsilateral or contralateral hemispheres. The shape of the burst-triggered potential differs between the VA and CM and is slightly affected by the distance. In line with the EEG results, the vigilance stage has a robust effect on the burst-triggered LFPs. The NREM evoked activity is much more pronounced. Notably, the difference between the effect of VA and CM bursts on LFP is much more prominent than in the previous analysis of their properties. In any case, the burst-triggered LFP and EEG indicate that the vigilance state robustly modulates thalamic-cortical dynamics. In the discussion below, we suggest this is why thalamic bursts do not awaken the cortex during NREM.

## Discussion

Rhythms of different temporal scales dominate the life of mammals, including humans. The awake-asleep circadian rhythm and the NREM-REM sleep ultradian rhythms are key examples of highly influential rhythms. Neural activity in the thalamus is correlated with, and probably plays a causal role in, the generation of these rhythms. Thalamic neurons burst during NREM sleep. It has been proposed that these bursts are periodic and synchronized^2,4^, thereby transmitting a null signal to the cortex^39^. Here, we recorded neuronal activity from the VA and CM thalamic nuclei, which receive BG GABAergic output. The discharge rate of BG output neurons decreases during NREM sleep^12^. This reduced BG output is expected to decrease the GABAergic drive to VA and CM neurons, leading to depolarization, elevated firing rates, and a shift toward tonic discharge. Nevertheless, we found that the discharge rate of VA and CM neurons decreased, and their tendency to discharge in bursts significantly increased during NREM sleep. Thus, the BG input to the VA and CM is probably a minor ^16,40,41^, and the activity of VA and CM neurons is more strongly affected by cortical^2^ or brainstem^19,20^ inputs. Quantitative analysis of the discharge patterns and synchronicity reveals that, contrary to the null hypothesis of sleep-synchronized periodic bursting, the thalamic bursts were neither periodic nor strongly synchronized. However, burst-triggered EEG and LFP responses varied across vigilance states, suggesting that NREM bursts do not wake the cortex due to altered dynamics of the thalamocortical network.

### *In vivo* thalamic bursts are characterized by a prolonged refractory period but lack clear periodicity

*In vitro* intracellular recording played a crucial role in understanding the role and mechanism of low-threshold calcium channels in the generation of thalamic bursts. However, brain slice preparations often exhibit more synchronous oscillations than those observed in the *in vivo* recordings^42,43^. Here, we recorded VA and CM spiking activity *in vivo* across natural vigilance states. Thalamic bursts were most common in the NREM stage. Notably, quantitative analysis of the burst discharge pattern of a large population of VA and CM neurons, in both the temporal and the frequency domains, reveals no significant evidence of periodic thalamic bursts (Fig. 2). Previous *in vivo* studies^6,7^ reporting the periodic behavior of thalamic bursts have not employed quantitative analysis. Visual inspection of their raw data figures aligns with our analysis.

### *In vivo* recorded bursts from both nearby and remote neuron pairs in the thalamus exhibit extremely weak correlation

As for periodicity, a synchronicity of neuronal activity is more commonly observed during low-arousal states and *in vitro* conditions. Physiological studies of thalamic brain slices^8,9^, as well as studies conducted under anesthesia^44^, often report synchronous activity. Our study of thalamic bursts reveals a broad central, low-amplitude, and symmetric peak (Fig. 3). The correlation intensity is maximal for close neuronal pairs during NREM sleep. It is minimal during REM sleep. In any case, our results indicate that the synchronicity of thalamic bursts during NREM is minor both in amplitude and time, suggesting that synchronicity is probably not the hallmark of the thalamic null message.

### Why do thalamic bursts not wake the cortex during NREM?

We first examined whether the thalamic bursts indeed fail to awaken the cortex—and, consequently, the animal. We found that thalamic bursts are not associated with the transition from sleep to the awake state, nor are they associated with eye opening, more eye movements, and increased muscular activity (Fig. 4a-f). Thus, the effects of thalamic bursts on cortical activity were further explored.

The frontal EEG and subcortical LFP were simultaneously recorded with the thalamic spiking activity. The EEG represents the summed activity of the underlying cortex. The subcortical LFP is a good proxy for whole-cortex and thalamic activity^37,38^. Our burst-triggered EEG and LFP (Fig. 4g, h) reveal clear and robust effects of the vigilance states. The NREM EEG and LFP activity exhibit an anticipatory negative deflection (indicating cellular depolarization) that begins one second before the thalamic burst. This probably suggests that during NREM sleep, cortical input is the primary driver of thalamic bursts. The post-burst EEG depolarization further suggests that the thalamic bursts are part of a closed thalamocortical loop that generates the delta rhythm^45^. In any case, our results indicate that the dynamics of the thalamocortical network during NREM sleep significantly differ from those during awake and REM sleep. We argue that these distinct dynamics underlie the varied effects of thalamic bursts on arousal and attention. An intriguing possibility is that this is due to shifts in cholinergic modulation of this network (Fig. 5a, b). The effects of thalamic bursts on the cortex during REM sleep closely resemble those observed during wakefulness. The brainstem atonia prevents the cortical awake-like activity from resulting in dream enhancement during REM sleep.

**Figure 5.**
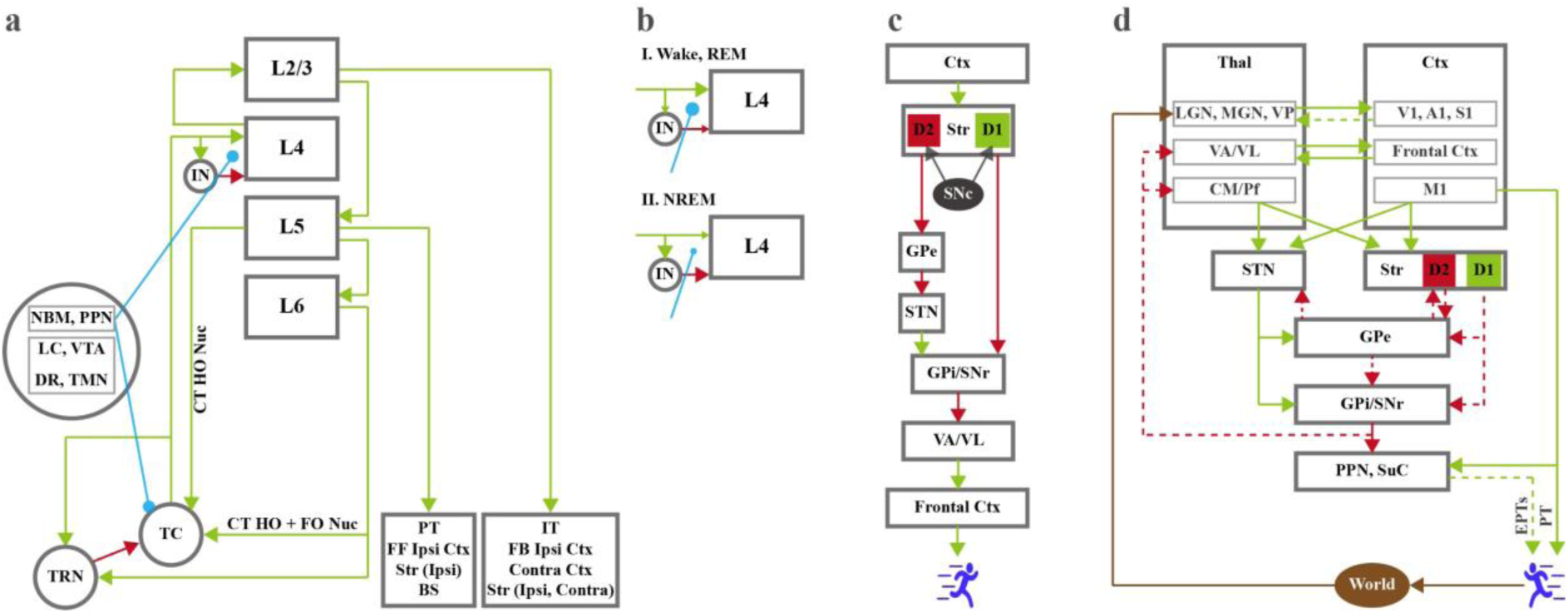
Schematic models of basal ganglia-thalamus-cortex network – revised. **a,** A schematic model of the thalamic cortical network. **b,** Suggested model of changes in cholinergic modulation of the cortex and their effects on cortical layer-4 connectivity during wake-sleep stages. **c,** The D1/D2 direct/indirect model of the basal ganglia-thalamus-cortex network. **d,** Revised model of the basal ganglia-thalamus-cortex modulatory/driving connectivity. Abbreviations. A1-primary auditory cortex; BS – brainstem; CM – Centromedian intra-laminar thalamic nucleus; CT HO Nuc– cortical thalamic tract to high-order thalamic nuclei; CT HO + FO Nuc – cortical thalamic tract to high- and first-order thalamic nuclei; D1/D2 – D1 and D2 striatal medium spiny neurons; DR – Dorsal Raphe (Serotonin); EPT – Extra-pyramidal tracts; FF – feed forward; FB – feedback; GPe, GPi – external and internal segments of the globus pallidus; IN -interneurons; IT – Intratelencephalic tracts (=> low-hierarchy cortex => striatum => contralateral cortex); L2/3 – cortex layer 2/3 (intra telencephalic neurons); L4 – cortex layer 4 (stellate neuron); L5 – cortex layer 5 (pyramidal and corticothalamic (to first order thalamic nuclei) neurons); L6 - cortex layer 6 (corticothalamic neuron (to first and higher-order thalamic nuclei)); LC – Locus coeruleus (Noradrenaline); LGN – lateral geniculate thalamic nucleus; M1 – primary motor cortex; MGN – medial geniculate thalamic nucleus; NBM – nucleus basalis of Mynert (Acetylcholine); PPN – Pedunculopontine nucleus (Acetylcholine); PT - pyramidal tract (=> spinal cord => brainstem => high-hierarchy Cortex); Pf - parafascicular intra-laminar thalamic nucleus; S1 – primary somatosensory cortex; SNc, SNr – substantia nigra compacta and reticulata; STN – subthalamic nucleus; Str – striatum; SuC – superior colliculus ; TMN – tuberomammillary nucleus (histamine); TC – Thalamocortical neuron; TRN –Thalamic reticular nucleus neuron; Thal – thalamus; V1 – primary visual cortex; VA – ventral anterior thalamic nucleus; VL-ventral lateral thalamic nucleus; VP – ventral posterior; VTA – Ventral tegmental area (Dopamine). Colour and shape coding: Light green – excitatory (e.g., glutamate) connection; Black red – inhibitory (e.g., GABA) connection; Black – dopaminergic connections; Cyan – cholinergic connection; Full and dashed lines, driving and modulatory connectivity; End arrow size reflects the connection efficacy.

### Is the BG-thalamus chain broken at night?

In 1989, Albin and colleagues published a groundbreaking model of the BG-thalamocortical network^10^. Over the years, BG researchers have proposed numerous modifications to this model^46^. Nevertheless, nearly all assumed that the BG output drives (inhibit or disinhibit) the thalamus and cortex^47,48^. Actions and movements are facilitated by reduced BG GABAergic flow in the thalamus, thalamic depolarization, and activation of the frontal cortex and descending motor pathways (Fig. 5c). Parkinson’s akinesia is due to the opposite changes in BG, thalamic, and frontal-cortex activity.

The researchers of the thalamus hold a different view. They classify the BG as modulators of thalamic activity. The current study suggests revising our BG models to align them with thalamic models^16,40,41^. The robust switch of VA and CM neurons to burst discharge during NREM does not align with the reduced discharge rate of their BG GABAergic afferents^12,23^. Notably, the effects of GABA on the membrane potentials (and, consequently, on discharge rate) are complex^41,47^. Further studies should directly test the impact of BG GABA on their thalamic targets. Currently, we suggest that the BG modulate the thalamus–frontal cortex (Fig. 5d). The functional connectivity between the BG and the thalamus may also be affected by the vigilance states. The BG may drive the thalamus in activated brain states, but modulate the thalamus during NREM sleep. Future studies may capitalize on the distinction between driving and modulation to investigate whether these mechanisms can be utilized to elucidate brain connectivity, including BG-to-brainstem connectivity, more effectively (Fig. 5d).

### Summary and Limitations

In this study, we recorded the activity of 1,161 VA and CM neurons across the natural cycles of vigilance states of two NHPs. The primary target was to explore the bursting activity of these neurons (1,254,441 bursts detected over 37 recording sessions). We found that VA and CM burst activity significantly increased during NREM sleep. However, these bursts were neither periodic nor tightly synchronized. Burst-triggered frontal EEG and thalamic LFP varied across vigilance states. We, therefore, conclude that state-dependent thalamocortical dynamics support different functional roles of thalamic bursts during activated brain states and NREM sleep, and that the BG modulate, but do not drive, their thalamic targets.

Our readers should be aware of the study’s limitations. The study is based on recordings from two NHPs. The results were consistent between these two NHPs. N = 2 is the standard practice in NHP research, and this is further justified by the 3R ethical rules for using animals in research. Nevertheless, further studies are needed. Similarly, our recordings are limited to only two thalamic nuclei out of more than twenty. Both nuclei are the targets of the BG. However, the CM is part of the intralaminar thalamic nuclei. Therefore, the similarity of VA and CM results might justify generalizing the heterogeneous collection of thalamic nuclei. Notably, the activity of the thalamic reticular nucleus, which plays a crucial role in regulating information flow in the thalamocortical network, has not yet been explored. Third, our data were simultaneously recorded only in left/right homologous structures, but not in serially connected structures (e.g., thalamus and cortex). A more comprehensive understanding of thalamocortical interactions requires simultaneous recordings from both structures and careful consideration of the immense diversity among cortical lamina and neurons^25,49^. Finally, BG GABA outflow during sleep was not measured. During NREM sleep, the discharge rate of GPi and SNr neurons decreased, accompanied by the emergence of a bursting discharge pattern. This increased burstiness may enhance GABAergic flow despite an overall reduction in firing rate. Further investigation using neurochemical methods can help clarify this effect^50^. We, therefore, hope that our results and conclusions will be supported by future multidisciplinary *in vivo*, *in vitro*, pharmacological, and physiological studies of different biological species, including humans.

## Author contributions

JG and HB conceived the research and designed the experiments. ZI, DW, and AR performed the surgical procedure. XL and JG supported the surgical procedure. They also performed the experiments, including electrophysiological and behavioral recordings, analyzed the data, and conducted the statistical analysis. XL, JG, and HB prepared the figures and wrote the manuscript. All authors read and approved the final manuscript. HB supervised the work.

## Acknowledgments

The authors would like to thank Uri Werner-Reiss, PhD, for his valuable support of the surgical procedures and all aspects of monkey care, Tamar Ravins Yaish, DMD, and the HUJI-ELSC animal facility team for their assistance. We thank Ad Aertsen for the fruitful discussion of correlation analysis and the association index, and Andy Horn, Jackie Schiller, Pnina Rapel, Aric Agmon, and Yuval Nir for their discussions and comments on early versions of the manuscript. We acknowledge the use of LLM tools in editing this manuscript.

## Data and code availability

Data and Matlab code will be available upon request from the corresponding authors. Please note that the code used in this study was developed by the researchers for data analysis and visualization. It is intended for research purposes and may not meet professional coding standards.

## Supplementary Information

1. Materials and Methods
2. Supplementary table and figures

## 1. Materials and Methods

### Animals

Data were obtained from two female vervet monkeys (Cercopithecus aethiops, monkeys Md and Wh) weighing 4-5 kg. Care and surgical procedures followed the National Research Council Guide for the Care and Use of Laboratory Animals^51^ and the Hebrew University guidelines for the care and use of animals in research. All experimental procedures were approved (MD-15-14412-5) and supervised by the Institutional Animal Care and Use Committee of the Hebrew University and Hadassah Medical Center, and the veterinary staff of the Hebrew University’s primate facility.

### Training and surgery

The non-human primates (NHPs) were habituated to sitting and sleeping in a primate chair within a dark, double-walled, sound-attenuating experimental room. They were trained to perform a modified memory-guided saccade using a video eye-tracker (ISCAN, 21 Cabot Road, Woburn, MA 01801, USA). After the initial training, the NHPs underwent four surgical procedures over a six-month period. In the first surgery, a head holder and two cranial (ground) screws were implanted, and the behavioral training (with a video eye-tracker) was conducted with the head fixed. In the next two surgeries, eye coils were implanted in both eyes^52^. Finally, a 34*27 mm craniectomy was done, and a recording chamber and frontal EEG skull screws were implanted in the skull in the fourth surgery. The surgeries were performed by a board-certified neurosurgeon (ZI), an ophthalmologist (DW), and an anaesthesiologist (AR), with the support of the research team (JG and XL), and under veterinary supervision. All surgeries were performed under general anaesthesia with appropriate antibiotics and pain relief medicine.

Following recovery from the last surgery, the precise location of the chamber was determined through a 3T MRI examination performed under moderate sedation (Medetomidine (Domitor) and Ketamine, i.m.). After the first (healthy condition) recording, the NHPs were treated with MPTP (1-methyl-4-phenyl-1,2,3,6-tetrahydropyridine), and the behavioral and neuronal activity were recorded in the Parkinsonian state. Here, we only report the results before the MPTP treatment. Upon completion of the experiment, all surgical attachments were removed from the NHPs. Monkey Md was then rehabilitated and placed at the Israeli Primate Sanctuary. Monkey Wh was euthanized to avoid suffering following discussions with the veterinary staff and the Institutional Animal Care and Use Committee.

### Recording Protocol

The NHPs were taken into the experimental room at around 4-5 PM during the week. They usually completed task performance within two hours and then slept for the entire night (from 7-8 PM until 4-5 AM, 5 nights per week), with the lights off but under infrared video and human supervision. They were food-restricted during the daytime and were fed during the task performance. Supplementary food was provided to the monkeys when they returned to the primate facility if their minimum daily caloric intake had not been met. They were housed in the monkey colony with their peers in a yard during the day and continuously on weekends.

During the recording sessions, the eye open/close state was tracked by an infrared eye video tracker (49-50 frame/s) and the eye position was continuously recorded with the bilateral eye-coil X-Y signals (7-SSCP-JGASM CUSTOM 13 ‘’COIL 13’’, 7-MTS-4340 3D / 4 sense coil, and 6-YET-H3 EYE COIL TEMPLATE 12 TO 20mm; Crist Instrument, Hagerstown, MD, USA). The EMG signal (trapezius muscle) and two frontal skull EEG signals were also continuously recorded. The eye-coils, EMG, and EEG signals were sampled at 2,750 Hz (SnR, Alpha Omega Engineering, Nof Hagalil, Israel).

The VA and CM thalamic nuclei were located based on the MRI examination (Fig. 1a), primate brain anatomic maps, and electrophysiological mapping^53^. During each recording session (night), up to four independently controlled microelectrodes were advanced separately into the targeted structures (VA or CM) in each hemisphere, for a total of up to eight electrodes per session. The microelectrodes were glass-coated tungsten, and their impedance ranged from 0.5 to 0.75 MΩ at 1000 Hz. Two experimenters (XL and JG) separately manipulate the Alpha Omega system (Electrode Positioning System, SnR, Alpha Omega Engineering, Nof Hagalil, Israel), each controlling four microelectrodes in one hemisphere. We only recorded the neuronal activity in homologous structures (e.g., left and right VA). The neural activity was hardware filtered by broadband 0.075 - 9000Hz 4-pole Butterworth hardware filters and continuously sampled at 44 kHz. The raw neural activity was online filtered to visually display (and store) LFP (0.075 – 300 Hz, sampling rate of 1,375 Hz) and SPK (300 – 9000 Hz, sampling rate of 44 kHz). The single-unit activity was detected and sorted online by manually setting an amplitude threshold and the shape of the action potential (template), as well as the maximal allowed deviation between threshold-crossing signals and the template. Up to four templates can be generated for each electrode. The data was synchronized and collected by AlphaLab SnR (Alpha-Omega Engineering, Nof Hagalil, Israel).

### Polysomnography analysis

A detailed description of our polysomnography methods can be found here^54^. In the current research, EMG was digitally offline bandpass-filtered in the 10 to 500 Hz range (stopband at 0 to 5 Hz, 520 to 1375 Hz). To minimize phase distortions, the forward-backward filtering was performed (this zero-phase filtering was used for any signal filtered offline). Sleep staging was performed using a semiautomatic staging algorithm that clustered 10-second nonoverlapping segments. Different vigilance stages (wakefulness, NREM, REM, and ambiguous/unclassified) were identified based on the eye-open fraction, the root mean square of the EMG signal, and the high/low EEG power ratio (the average power at 15 to 25 Hz / the average power at 0.1 to 7 Hz). The segment would be classified as NA (Not Available) if the missed length of any signal (Eye open state, EEG, EMG, Coil eye-position, LFP, SPK) was longer than 11*(1/SR)*1000 ms (out of 10,000 ms); SR is the sampling rate of the corresponding signal. Before semiautomatic clustering, 10% of the night epochs were scored manually by a trained expert (JG). Both left and right EMGs were used separately for the staging analysis. The better staging results provided by the semiautomatic algorithm, which matched the expert staging in more than 85% of the tested segments, were accepted for further analysis.

Eye-coil signals were converted from voltage to degrees (angular position) using calibration data obtained on the same day. The transform relationship was generated using the fit geometric transformation^55^. The ‘projective’ transformation type was used since the original calibration data demonstrated that the scene appeared tilted. This transformation maintained the straight lines straightness and converged parallel lines toward a vanishing point. The converted eye-coil signals were used to calculate the velocity and acceleration of the eyes, as well as to identify saccades and drifts. Only saccades/drifts with a magnitude larger than one degree were kept for further analysis.

The basic sleep staging was refined based on the eye open percentage, EMG, and the saccade frequency of the right eye. Typically, the EMG RMS is relatively larger in NREM sleep and smallest in REM sleep (Fig. 1b, d and Fig. 4f). Therefore, if a REM segment exhibits an EMG RMS value larger than the average EMG RMS of the NREM segments, or if an NREM segment shows an EMG RMS value smaller than the average EMG RMS of the REM segments, these segments will be considered unreasonable. This was done for both left and right EMG. The left/right EMG with relatively less unreasonable segments would be kept for further analysis. The NREM segments that occurred near REM epochs (two segments before and five after the previous REM segment) and additionally were characterized with eye-open ratio close to zero, more than 0.5/s eye saccades, and low EMG activity (the left/right EMG not larger than two times of the maximum EMG RMS during REM) were identified as REM candidates. For each candidate REM segment, if the duration of REM within the surrounding segments (three segments before and after the REM candidate) was longer than that of not-REM segments (NREM, wakefulness, or unclassified), this candidate REM would be classified as REM; otherwise, it would be identified as NREM. The awake or unclassified segments demonstrating an eye-open fraction between 0.4 and 0.7 were identified as wakeful candidates. These candidates would be classified as awake or unclassified segments using a method similar to the one used to determine the REM candidates. Finally, in the original classification of sleep stages, the eye open fraction must be larger than 0.6, 0.4-0.6, and smaller than 0.4 in the awake, unclassified, and sleep (NREM and REM) states, respectively. The NREM segments with eye-open fractions between 0.05 and 0.3 would be refined as unclassified if the length of unclassified time within the surroundings of this NREM candidate was longer than the NREM sleep time. An example of a one-night hypnogram and PSG example is shown in Fig. 1b.

### Spike Analysis

Spiking (300-9000 Hz) signals were filtered using an IIR comb notch filter with 881 (44,000/50 + 1) notches, which removed the 50 Hz power noise and its harmonics from the signal, given a sampling rate of 44,000 Hz. The Q factor for this filter was set to 35, namely, a highly sharp notch filter. The filtered spiking activities were rectified by absolute value, and the mean of the rectified vector was subtracted to obtain multi-unit activities (MUA). A low-pass Butterworth filter (210 Hz passband frequency, 260 Hz stopband frequency, 1 dB passband ripple, and 5 dB stopband attenuation) was designed to filter the MUA. The filtered MUA was down-sampled to 1/32 of the original sampling rate (yielding a 1,375 Hz sampling rate) by averaging the amplitude of the surrounding 32 sampling points. The filtered and down-sampled MUA was used to calculate its auto- and cross-correlation histograms after being segmented into 10-second epochs corresponding to the sleep staging epochs. The mean value of this 10-second MUA segment was subtracted to minimize the DC (zero frequency) power. The lag range of the auto- and cross-correlation histograms was from -2 to 2 seconds. The auto- and cross-correlation histograms were calculated using the built-in function of MATLAB 2020b (xcorr, the correlation coefficient method), and smoothed by convolution with a Gaussian kernel having a standard deviation of 3.6 ms and 43.6 ms (5 and 60 times the time resolution, i.e., 1/1375 s), respectively. The xcorr correlation coefficient method ensures that the correlation coefficients (R) values could range from -1 to 1.

The action potentials (spikes) of well-isolated neurons (Isolation Score > 0.7 or > 0.85)^24^ were transformed into a continuous binary train of 0/1 values, representing single-unit activity (SUA). The SUA was down-sampled to 1/32 of the original sampling rate (i.e., from 44 kHz to 1,375 Hz) by summation of its trains. The spike train of every neuron was also separated into 10-second epochs based on the segments of sleep stages. The firing rate (FR, the number of spikes/s), the inter-spike intervals (ISI), and the coefficient of variation of the ISIs (CV-ISI, defined as STD(ISI)/Mean(ISI)) were calculated for every 10-second segment, and then were clustered into three groups (wakefulness, NREM, and REM) based on the awake-sleep staging. For each neuron, the FR, ISI, and CV-ISI were averaged within each group.

The power spectrum density (PSD) of SUA was calculated using a 10-second moving window, with a 5-second moving step, a frequency range of 0.1 Hz to 100 Hz, and a frequency resolution of 0.1 Hz. For each 10-second binary (0/1) train, we subtracted the mean to minimize the DC (Frequency = 0 Hz) power. The PSD unit is therefore given as NormSpk^2^/Hz.

The auto- and cross-correlation (R-values) histograms of spike trains were calculated using the built-in function of MATLAB 2020b (xcorr, the correlation coefficient method). For this method, the mean value of each 10-second spike train was subtracted to show the negative and positive correlation values. The lag range and normalization were the same as for the MUA correlation analysis.

The conditional discharge rate correlation histograms were calculated as the number of spikes (spike count) of the reference (triggered) cell, normalized by the number of spikes of the trigger neuron and the bin duration. The lag range was also from -2 to 2 seconds. Edge effects were corrected by normalizing with the actual number of valid trigger spikes and bin duration. For example, at specific lag time points, a triggered spike train may be absent when aligned to a trigger spike, resulting in one fewer valid trigger spike contributing to that time point. The valid trigger count was adjusted by subtracting one from the total number of trigger spikes in such cases.

The results from the conditional discharge method were used to calculate the association index (AI), which indicates the fraction of spikes contributed by the common-input mechanism out of total spikes. AI can be calculated as the area of the original cross-correlation histogram (counts/bin) peak divided by the total number of spikes from the triggered neurons. We used two ways to calculate the AI:

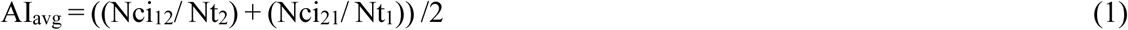

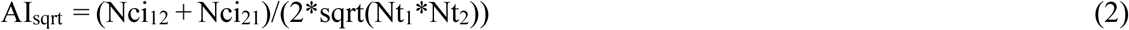

AI_sqrt_ /AI_avg_ is the fraction of the spikes of the two studied neurons (1 and 2) generated by the common (shared) input to the neurons. Nci_12_ indicates the number of spikes in the central (common input) peak (neuron 1 is the trigger neuron, and neuron 2 is the triggered one). Nci_21_ indicates the number of spikes in the central peak, but the trigger/triggered neurons are switched. Nt_1_ or Nt_2_ is the total number of spikes of the triggered neurons. The base for calculating the Nci is the average firing rate of the triggered neuron, and the Nci peak range is from -0.5s to 0.5s. Neurons 1 and 2 were referred to as two neuron pairs with repetition or one neuron pair without repetition. In the ‘with repetition’ condition, the pair was treated directionally, such that neuron 1 triggering neuron 2, and neuron 2 triggering neuron 1, were considered as two distinct pairs; in the ‘without repetition’ condition, both directions were used for the same calculation, and thus counted as a single neuron pair.

Before getting the R values at time 0 and calculating the AI, the cross-correlation histograms were smoothed by Gaussian convolution with 43.6 ms (60 times the time resolution (1/1375 s)) standard deviation. The same smoothing method was also applied to the autocorrelation histograms and PSD. The standard deviations of the smoothing kernel for the autocorrelation histograms and PSD smoothing are 0.36 ms (0.5*(1/1375)*1000) and 0.2 Hz (2 times the frequency resolution (0.1 Hz)), respectively. Such light smoothing was applied to auto-correlation to minimize the strong effect of maximal values (1 for normalized autocorrelation histograms) at time zero.

### Burst detection

Bursts were detected using the maximal interval method (MI, https://www.neuroexplorer.com/downloads/NeuroExplorerManual.pdf)^28^ and the Poisson surprise (PS) method^29^. In the MI method, we set the maximum length of the first inter-spike interval (ISI) of a candidate burst to be 10 ms. We added spikes until we reached the maximum ISI limit (12 ms). A burst should display a minimum silent period of 50 ms (with zero or no more than one spike) before the burst. Every burst should include at least three spikes.

In the PS method, we identified the burst candidate as at least three consecutive spikes with ISIs shorter (each one of them) than 1/10 of the average ISI in a 10-second epoch (ISI limit). Spikes are added to the end of the candidate burst until the ISI of the added spike is larger than 1.5 times the ISI limit or the number of added spikes is 5. We then calculated the PS of the candidate burst as a basic reference. Spikes will be removed from the beginning of the candidate burst if this maximizes the PS value. The removal process will be repeated until five spikes are removed, or the PS value is maximized. Ultimately, a candidate burst would be considered as a valid burst only if its PS value was at least 10.

### Burst analysis

Burst features, i.e., burst frequency, inter-burst interval (IBI), the coefficient of variation of IBI (CV-IBI), burst duration, the number of spikes per burst, the intra-burst firing rate, and the burst ISI ordinal duration, were analysed. Burst frequency indicates the number of bursts per second. IBI is the duration between the beginning (first spike) of the current (n) burst and the beginning (first spike) of the next (n + 1) burst. Burst durations were calculated as the duration between the burst’s first and last spikes, and the typical duration of a single spike (96 sampling points) was added. The number of spikes per burst was divided by its corresponding burst duration to yield the intra-burst firing rate.

The burst spikes were removed from the SUA to get the tonic activity. The FR, ISI, and CV-ISI were calculated for tonic activity. The same down-sampling features as used for the SUA were applied to tonic activity and to burst trains, where each burst was represented as a binary (0/1) event occurring at the time of the first spike in the burst. We also applied the same auto- and cross-correlation analyses described above for single-unit activity to tonic and burst activity. The cross-correlation histogram of burst trains in a particular vigilance state (e.g., NREM sleep) calculated by the conditional discharge method were removed from the analysis database if the trigger or triggered neuron had only one burst/10 s, both trigger and triggered neurons had bursts only near the same 2-second edge of the 10-second burst trains, and only one segment was available in this sleep stage. A total of 67 burst pairs without repetition were removed from the database of 4,416 burst pairs without repetition. The auto- and cross-correlation histograms of tonic activity were smoothed by convolution with a Gaussian kernel having a standard deviation of 0.36 ms (0.5*(1/1375)*1000) and 43.6 ms (60*(1/1375)*1000), respectively. For the burst trains, the standard deviations of the Gaussian kernel were 14.5 ms (20*(1/1375)*1000) for the auto-correlation histograms, and 43.6ms (60*(1/1375)*1000) for cross-correlation histograms. The auto-correlation of burst trains was smoothed relatively stronger than that of SUA and tonic activity, because it had a longer refractory period. The same methods for calculating the AI and PSD of spike trains were also applied to tonic activity and burst trains. Therefore, the units of the PSDs for tonic activity and burst train are NormSpk^2^/Hz and NormBst^2^/Hz, respectively. The PSD of SUA, tonic activity, and burst trains was analyzed over the 0.1 - 10 Hz frequency range. Additionally, cross-correlation (R-values) histograms of spike trains, burst trains, and tonic activity with 1-second time resolution were also calculated using the correlation coefficient method (xcorr, the built-in function of MATLAB 2020b, Fig. S8).

The percentage of spikes within bursts among all spikes in the 10-second epoch is defined as the probability of burst spikes. The firing mode of every 10-second segment was identified based on the probability of spikes occurring in bursts (burst spikes). Therefore, three firing modes were defined: tonic (< 3%), mixed (3%-30%), and burst (≥30%) mode.

### Analysis of the relations of bursts with behavioral and other physiological metrics

The burst frequency of every 10-second epoch, the probability of burst spikes, and the number of spikes per burst were aligned with the transition of sleep stages (e.g., transitioning from NREM to REM). To overcome the confounding effects of the inherently different percentage of vigilance stages, the corresponding predicted number of bursts was calculated using the average burst frequency of each neuron in wakefulness, NREM, or REM, and the percentage of the three sleep stages:

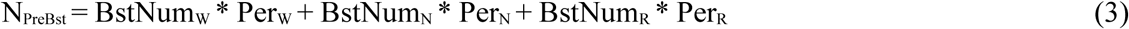

N_PreBst_ is the predicted number of bursts. BstNum_W_, BstNum_N_, and BstNum_R_ indicate the average number of bursts, the probability of burst spikes or the number of spikes per burst in the wakefulness, NREM, and REM, respectively. Per_W_, Per_N_, and Per_R_ indicate the percentage of the three vigilance states, respectively. The difference between the real and the corresponding predicted number of bursts was calculated.

The two firing modes (burst and tonic) were combined with the three vigilance states (wakefulness, NREM, and REM). Therefore, there are six mode-stage combinations (burst-wakefulness, tonic-wakefulness, burst-NREM, tonic-NREM, burst-REM, and tonic-REM).

We aligned the sleep stages to these six mode-stage combinations to explore the relationship between sleep states and firing mode. Only neurons having both tonic and burst firing mode segments were included for further analysis. Neurons having fewer than three aligned segments of wake-sleep stages were excluded. The difference between the percentage of the sleep stage related to the burst and tonic firing mode segments was calculated. Therefore, positive values indicated that the burst boosts staying in the same wake-sleep state or switching to another state.

The eye-open/close state, rapid eye movements (saccades) frequency, slow eye movement (drifts) frequency, and EMG were aligned to bursts to reveal fast changes, which might be masked by the 10-second temporal resolution of our behavioral analysis. The eye open-close state is represented by the percentage of eye closure, i.e., the number of eye closures per second divided by the average frame rate of the video. A 4^th^-order bandpass Butterworth filter with a cut-off frequency of 8 Hz to 750 Hz was used to obtain the 10 - 500 Hz EMG signal. This filtered EMG was down-sampled from 2,750 Hz to 1,375 Hz by averaging its amplitude. The absolute value of the EMG (rectification) was calculated before averaging the filtered, down-sampled, and burst-aligned EMG. The burst-aligned eye open-close state, saccade frequency, and eye-drift frequency were smoothed by convolution with a Gaussian kernel with 50 ms (50*(1/1000)*1000) standard deviation. A Gaussian kernel with a 36.4 ms (50*(1/1375)*1000) standard deviation was used for the EMG.

The frontal EEG and thalamic LFP (VA and CM) were also aligned to the bursts to study the relationship between the thalamic bursts and these extra- and intra-cranial physiological signals. The 50Hz power artifact and its harmonics (100Hz, 150Hz, 200Hz, and 250Hz) of EEG and LFP were removed by a second-order IIR notch filter. A low-pass Butterworth filter (210 Hz passband frequency, 260 Hz stopband frequency, 1 dB passband ripple, and 5 dB stopband attenuation) was also designed to filter the EEG and LFP. The EEG signals were resampled as the EMG signal. For EEG and LFP, the mean value was subtracted from the filtered and down-sampled signals before alignment to remove the DC component of the signal. We used a convolution with a Gaussian kernel having a standard deviation of 36.4 ms (50*(1/1375)*1000) to smooth the EEG and LFP signals. Finally, they were normalized by z-score based on their baseline (from two to one second before the burst).

### Statistical analysis

Statistical analysis was performed using MATLAB R2020b. The p-value was calculated using the Wilcoxon rank sum test for the independent data and the Wilcoxon signed-rank test for the two matched samples. The statistical tests were two-tailed. A P-value threshold of 0.05 was used, and the results were corrected for multiple comparisons using the Bonferroni method.

## 2. Supplementary table and figures

**Table S1.** The neuronal database.

| | | Isolation score $\geq 0.7$ | | | Isolation score $\geq 0.85$ | | |
| --- | --- | --- | --- | --- | --- | --- | --- |
|  | NHP/Structure | VA | CM | VA+CM | VA | CM | VA+CM |
| Nights | Md | 11 | 8 | 19 | 11 | 7 | 18 |
|  | Wh | 9 | 9 | 18 | 9 | 9 | 18 |
|  | Md+Wh | 20 | 17 | 37 | 20 | 16 | 36 |
| Neurons | Md | 326 | 178 | 504 | 220 | 100 | 320 |
|  | Wh | 363 | 294 | 657 | 176 | 164 | 340 |
|  | Md+Wh | 689 | 472 | 1161 | 396 | 264 | 660 |
| Bursts<br>(MI<br>detection<br>method) | Md | 198,347 | 184,701 | 383,048 | 135,434 | 111,392 | 246,826 |
|  | Wh | 369,978 | 501,415 | 871,393 | 167,352 | 284,744 | 452,096 |
|  | Md+Wh | 568,325 | 686,116 | 1,254,441 | 302,786 | 396,136 | 698,922 |

Number of recording sessions (nights), well isolated (isolation quality $> 0.7, 0.85$ ) recorded neurons and detected (using the MI method) bursts in the ventral anterior and centromedian (VA and CM) thalamic nuclei of the two NHPs (Md and Wh) that participated in the study. Abbreviations. NHP – non-human primate; MI – maximum interval burst detection method.

**Figure S1.**
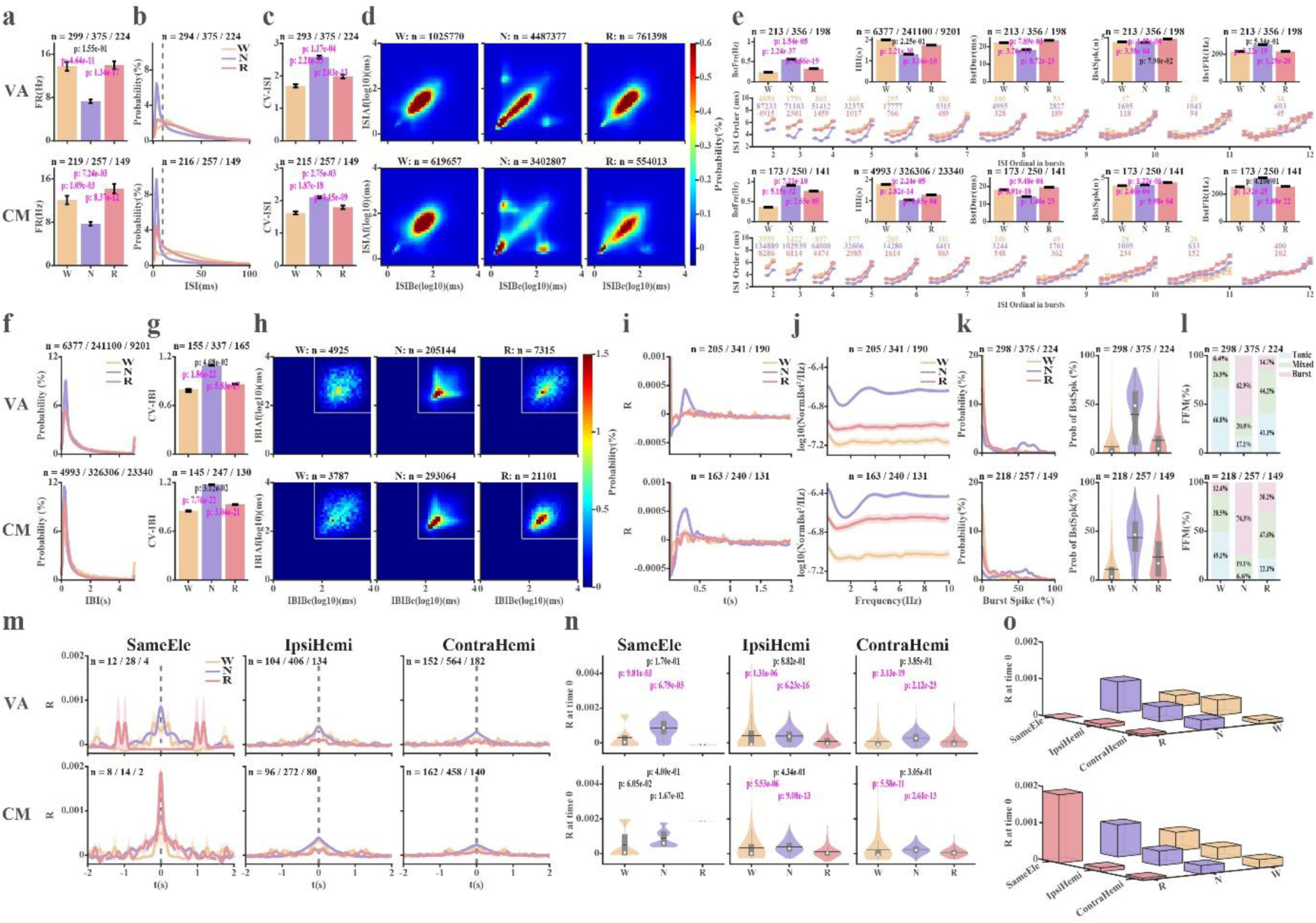
Discharge properties of spikes and bursts in thalamic neurons with Isolation quality ≥ 0.85. **a**-**d**. spike properties. **e**-**o**. Burst properties. **a**, Firing rate. **b**, Inter spike interval (ISI) histogram. The vertical dashed lines are located at 10 ms. **c**, Coefficient of variation of the ISIs (CV-ISI). **d**, ISI return map of ISI(n+1) (ISIAf) versus ISI(n) (ISIBe). **e**, Burst features. Row 1^st^, from left to right: Burst frequency (BstFre), Inter-burst interval (IBI), burst duration (BstDur), number of spikes in bursts (BstSpk), intra-burst spike discharge rate (BstFR). Row 2^nd^, ordinal inter-spike intervals in bursts of different lengths. Only bursts with fewer than 12 intervals are shown. **f**, Inter-burst interval (IBI) histogram. **g**, CV-IBI. **h**, IBI return map (IBI(n+1) (IBIAf) vs. IBI(n) (IBIBe)). Fifty milliseconds are the minimum silent time before a burst in the maximum interval burst detection method. Therefore, there are no IBIs below and to the left of the horizontal and vertical white lines (log10(50) ms). The dark blue pixels in this region indicate NaN (Not a Number). **i**, Auto-correlation histograms of burst trains. **j**, Power spectrum densities of burst trains. **k**, Fraction of spikes within bursts out of total spikes (left), and distribution of the burst spikes fraction (right). **l**, The percentage of 10-second segments with burst, mixed, and tonic firing modes. **m**, Cross-correlation histograms of burst trains simultaneously recorded from the same electrode, the ipsilateral hemisphere or contralateral hemisphere (SameEle, IpsiHemi, and ContraHemi). **n**, The distribution of R values at time 0 in **m**. The number of R values at time 0 is the same as that of neuron pairs with repetition in **m. o**, 3D-bar plots of the average R values at time zero as a function of wake-sleep state and distance. Note: n = 2 neuron-pairs with repetition is the outlier of CM R value at time 0 recorded from the same electrode during REM sleep. In all subplots: Up-VA, bottom-CM. Abbreviations. W-Wakefulness, N-NREM, R-REM. Colour coding: Yellow-Wakefulness, Purple-NREM, Pink-REM. Data are shown as average ± SEM (standard error of the mean) in **a**, **c**, **e**, **g**, **i**, **j**, and **m**. The n indicates the number of ISI pairs in **d**, IBIs in subplot **e**-2^nd^ column and **f**, IBI pairs in **h**, bursts in 2^nd^ and 4^th^ rows of **e**, neuron pairs with repetition in **m**, and neurons in the rest of the subplots. A Bonferroni-corrected Wilcoxon rank sum test was used to calculate the statistical significance of the difference in the FR (**a**), CV-ISI (**c**), burst features (**e** and **g**), and R values at time zero (**n**) between different vigilance stages (p < 0.05/3). P values marked in magenta indicate a significant difference; otherwise, they are black. Related to Figures 1, 2, and 3.

**Figure S2.**
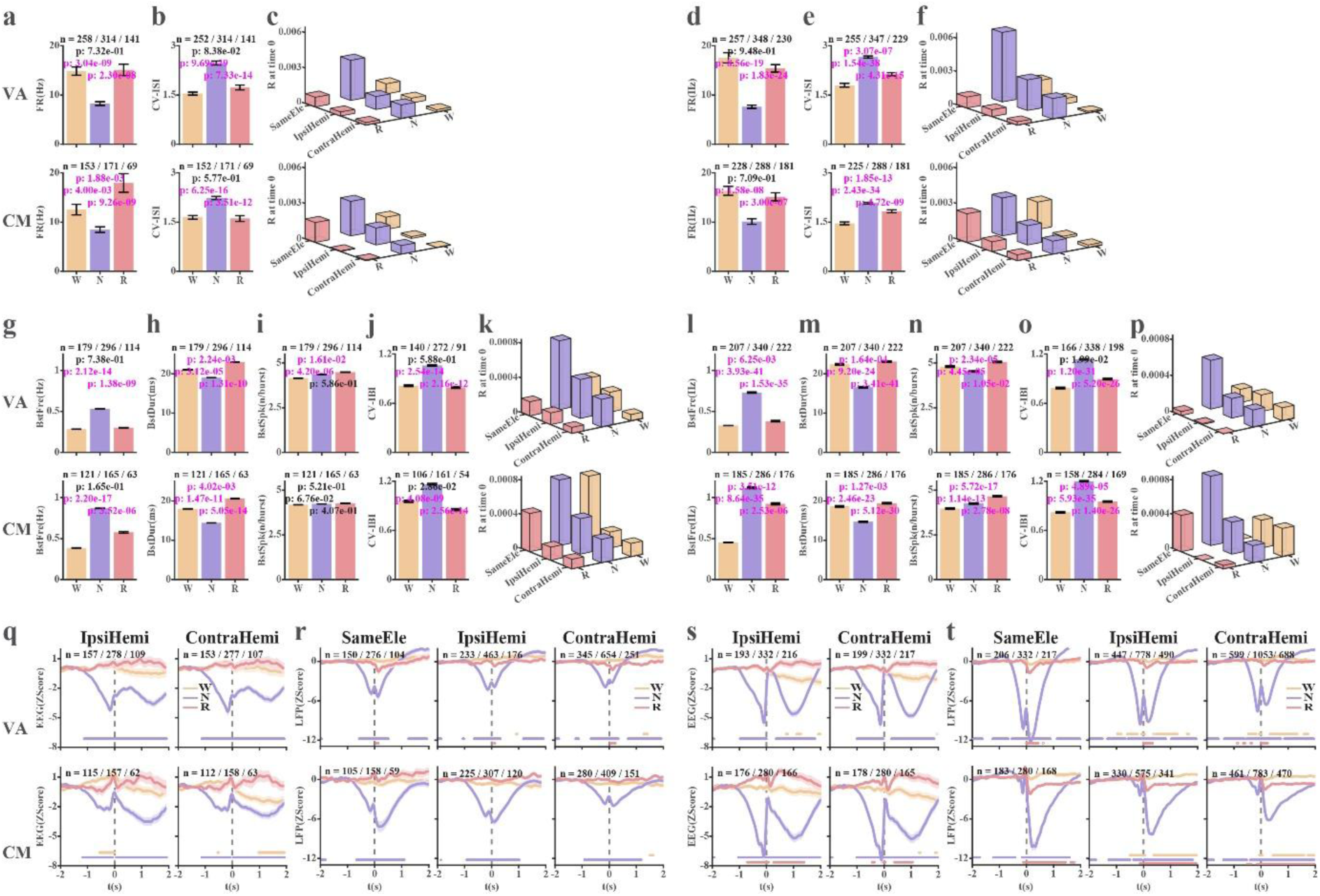
Neuron activity of two NHPs (Monkey Md and Wh). The left panel shows the results of Md, and the right panel shows the results of Wh. **a** and **d**, Firing rate (FR) of single unit activity (SUA). **b** and **e**, Coefficient of variation of the inter-spike interval (CV-ISI) of SUA. **c** and **f**, 3D-bar plots of cross-correlation R values at time zero (bin width = 0.7ms) of SUA, presented as a function of vigilance stages and distance. **g** and **l**, Burst frequency (BstFre). **h** and **m**, Burst duration (BstDur). **i** and **n**, The average number of spikes per burst (BstSpk). **j** and **o**, Coefficient of variation of the inter-burst interval (CV-IBI). **k** and **p**, 3D-bar plots of cross-correlation R values at time zero (bin width = 0.7ms) of burst trains, presented as a function of vigilance states and distance. **q** and **s**, Burst-triggered frontal EEG from the ipsilateral or contralateral hemisphere. **r** and **t**, Burst triggered thalamic LFP simultaneously recorded from the same electrode, or different electrodes in the ipsilateral or contralateral hemisphere. Data is shown in mean ± SEM, and n indicates the number of neurons in **a**, **b**, **d, e**, **g**-**j**, **l**-**o**, and **q**-**t**. A Bonferroni-corrected Wilcoxon rank sum test was used to calculate the statistically significant difference. P-value indicating a statistically significant difference is marked in magenta; otherwise, it is marked in black in **a**, **b**, **d, e**, **g**-**j**, and **l**-**o** (p < 0.05/3). In **q**-**t**, the colored circles show the statistically significant difference (p < 0.05/1375). Abbreviations and color coding as in Fig. S1. Related to Figs 1**g, h** and **n**, 2**b** and **d**, 3**d**, and 4**g** and **h**.

**Figure S3.**
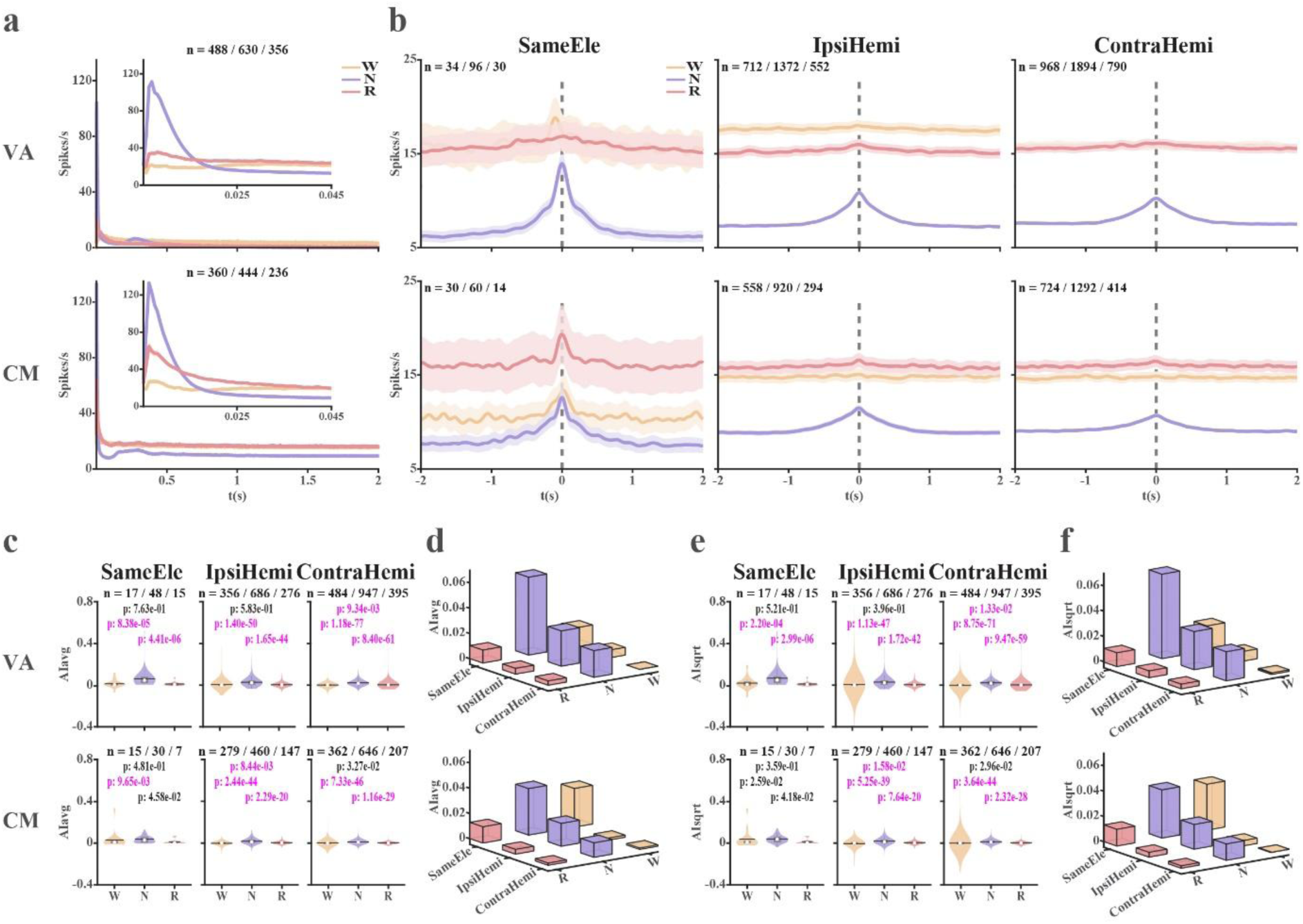
Auto- and cross-correlation conditional discharge rate histograms of thalamic single unit activity. **a**, Auto-correlation histograms of spike trains (lag = 2 s). The insets provide a magnified view of the auto-correlation histograms at a lag 50 ms. **b**, Cross-correlation histograms of spike trains. **c**, Violin plots of the distribution of Association Indices (AI, fraction of spike train contributed by common-input mechanism) calculated as AI_avg_ (equation 1). **d**, 3D-bar plots of average AI_avg_ values as a function of the vigilance stage and distance. **e**, AI calculated as AI_sqrt_ (equation 2). Violin plots of the distribution of AI_sqrt_ values, **f**, 3D-bar plots of average AI_sqrt_ values as a function of the vigilance stage and distance. Data in **a** and **b** are shown in mean ± SEM. The n indicates the number of neurons in **a**, neuron pairs with repetition in **b**, and the neuron pairs without repetition in **c** and **e**. The Wilcoxon rank sum test was used to calculate the statistically significant differences of AI (**c** and **e**) among different vigilance stages. The statistically significant difference was corrected by the Bonferroni method (p < 0.05/3). The p-values indicating a significant difference are marked in magenta. Otherwise, they are in black. Abbreviations and colour coding as in Fig. S1. Related to Fig. 1.

**Figure S4.**
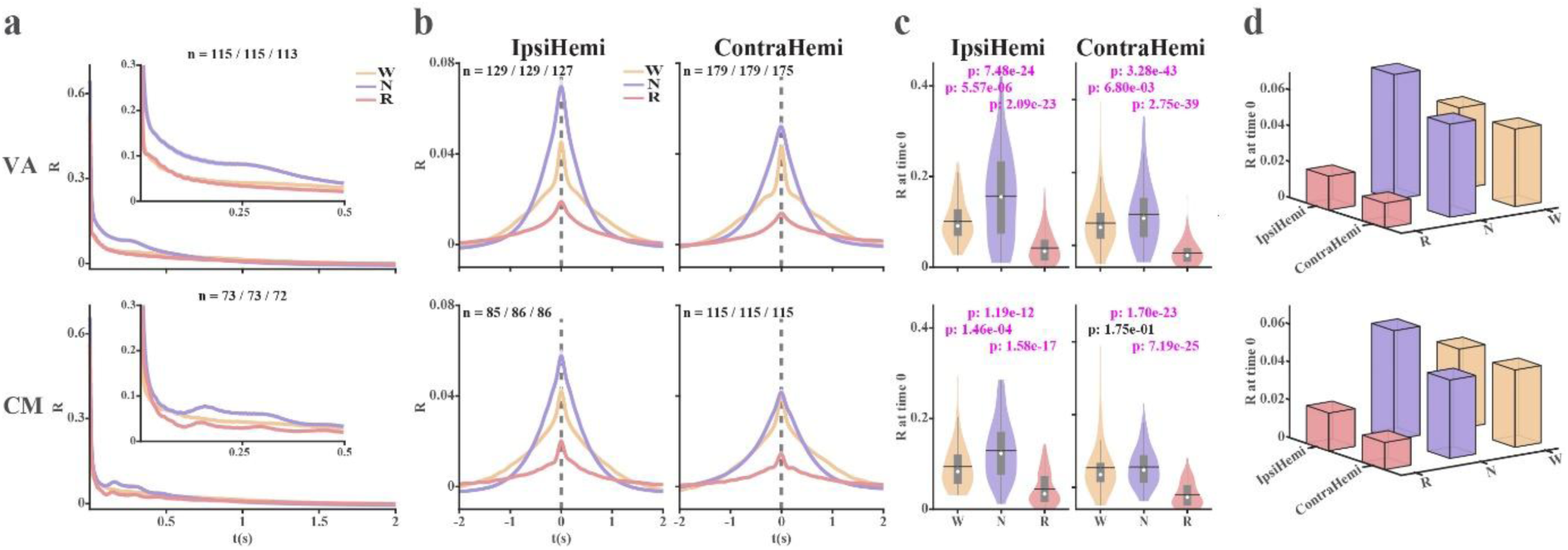
Auto- and cross-correlation histograms of thalamic multi-unit activity (SPK-MUA). Spiking activity (300 - 6000 Hz, SPK is rectified by the absolute operator, and its mean is subtracted, to represent the multi-unit-activity, MUA). **a**, Auto-correlation histograms of the spiking activity. The insets provide a magnified view of the auto-correlation histograms at a lag of 0.5 s. **b**, Cross-correlation histograms of the MUA. **c**, The distribution of R values at time zero. The number of R values at time zero is the same as the number of electrode pairs, as shown in **b**. **d**, 3D-bar plots of average R values as a function of vigilance state and distance. The broadband spiking activity measures the summated activity of all units near the electrode; therefore, it does not enable correlation analysis of different units recorded by the same electrode. Data in **a** and **b** are shown in mean ± SEM. The n means the number of electrodes in **a** and the number of electrode pairs in **b**. The Wilcoxon rank sum test was used to calculate the statistically significant difference of R values at time zero, which was corrected using the Bonferroni method, p < 0.05/3. P values marked in magenta show a statistically significant difference; otherwise, in black. Abbreviations and colour coding as in Fig. S1. Related to Fig. 1.

**Figure S5.**
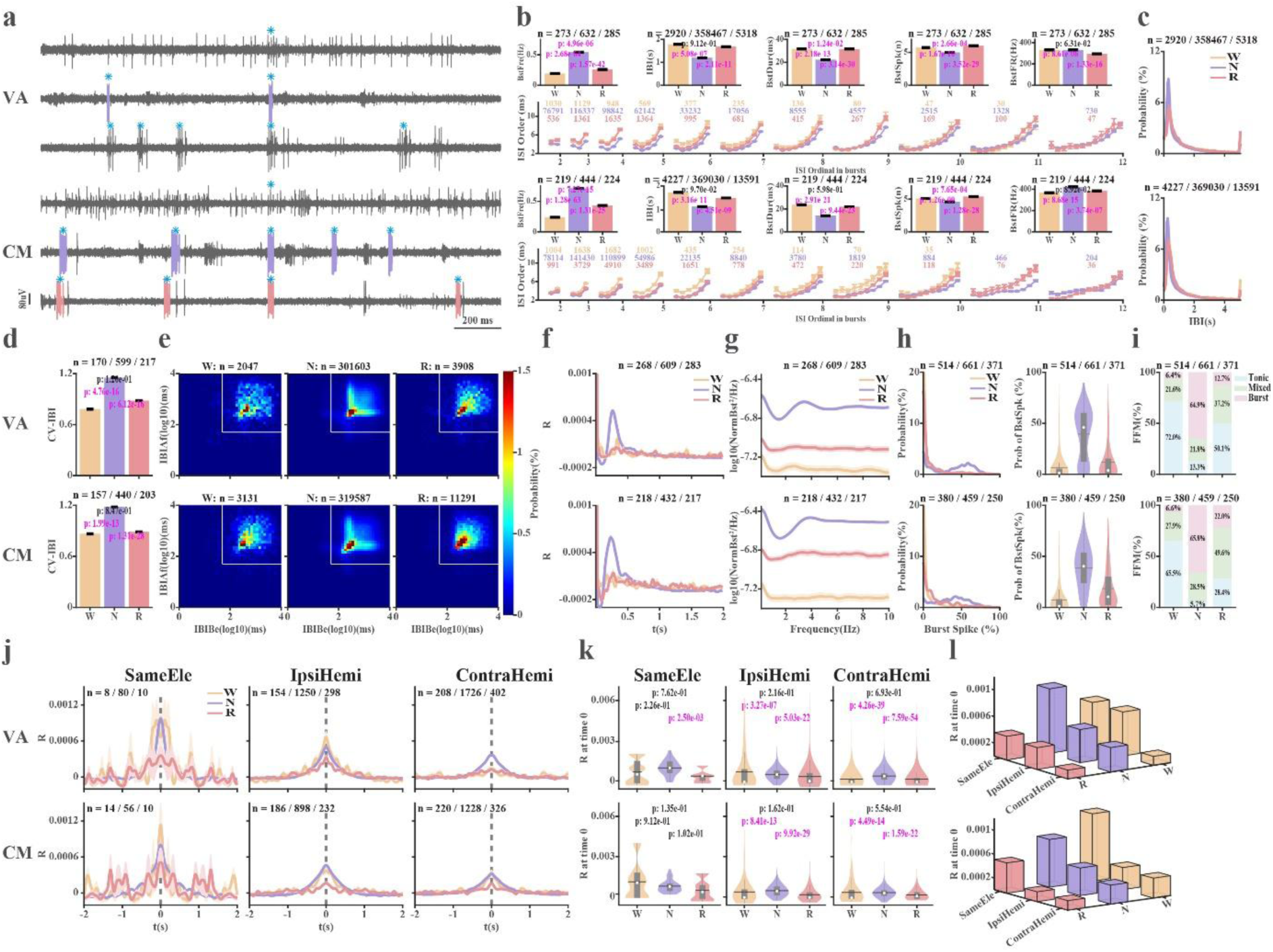
Burst properties analysis for bursts detected by the Poisson Surprise method. Bursts were detected from VA and CM neurons with an isolation quality larger than 0.7 using the Poisson Surprise (PS) method. **a**, the same examples as shown in Figure 2. Bursts marked in color (color code of sleep stages) are detected by the PS method, marked in cyan stars are bursts detected by MI methods. From top to bottom, the examples are from W, N, and R in VA, and CM. **b**, Burst features. Row 1^st^: Burst frequency (BstFre), Inter-burst interval (IBI), burst duration (BstDur), number of spikes in bursts (BstSpk), and intra-burst discharge rate (BstFR). Row 2^nd^, ordinal inter-spike intervals in bursts of different lengths. Only bursts with fewer than 12 intervals are shown. **c**, IBI histogram. **d**, CV-IBI. **e**, IBI return map (IBI(n+1) (IBIAf) vs IBI(n) (IBIBe)). The white line indicates log_10_(50) ms for comparison with the IBI return map of the burst detected by the maximum interval method (Fig. 2**e** and Fig. S1**h**). Notably, there is no limitation on silent time before a burst for the PS method. **f**, Auto-correlation histograms of burst trains. **g**, PSD of burst trains. **h**, Fraction of burst spikes out of total spikes (left), and distribution of fraction of burst spikes out of total spikes (right). **i**, Percentage of burst/mixed/ tonic firing modes across vigilance stages. **j**, Cross-correlation histograms of burst trains of neuronal pairs recorded from the same electrode, the ipsilateral hemisphere, or contralateral hemisphere (SameEle, IpsiHemi, and ContraHemi). **k**, violin plots of the distribution of R values at time 0 in **j**. The number of R values at time 0 is the same as that of neuron pairs with repetition shown on the cross-correlation histogram (**j**). **l**, 3D-bar plots of the average R values at time zero as a function of vigilance state and distance. Data are shown as average ± SEM in **b**, **d**, **f**, **g**, and **j**. The n indicates the number of IBIs in subplot **b**-2^nd^ column and **c**, IBI pairs in **e**, bursts in 2^nd^ and 4^th^ rows of **b**, neuron pairs with repetition in **j**, and neurons in the rest of the subplots. A Bonferroni-corrected Wilcoxon rank sum test was used to calculate the statistically significant differences in burst features (**b** and **d**) and R values at time 0 (**k**) among different vigilance stages (p < 0.05/3). P values marked in magenta indicate a significant difference; otherwise, they are marked in black. Abbreviations and colour coding as in Fig. S1. Related to Figs. 2 and 3.

**Figure S6.**
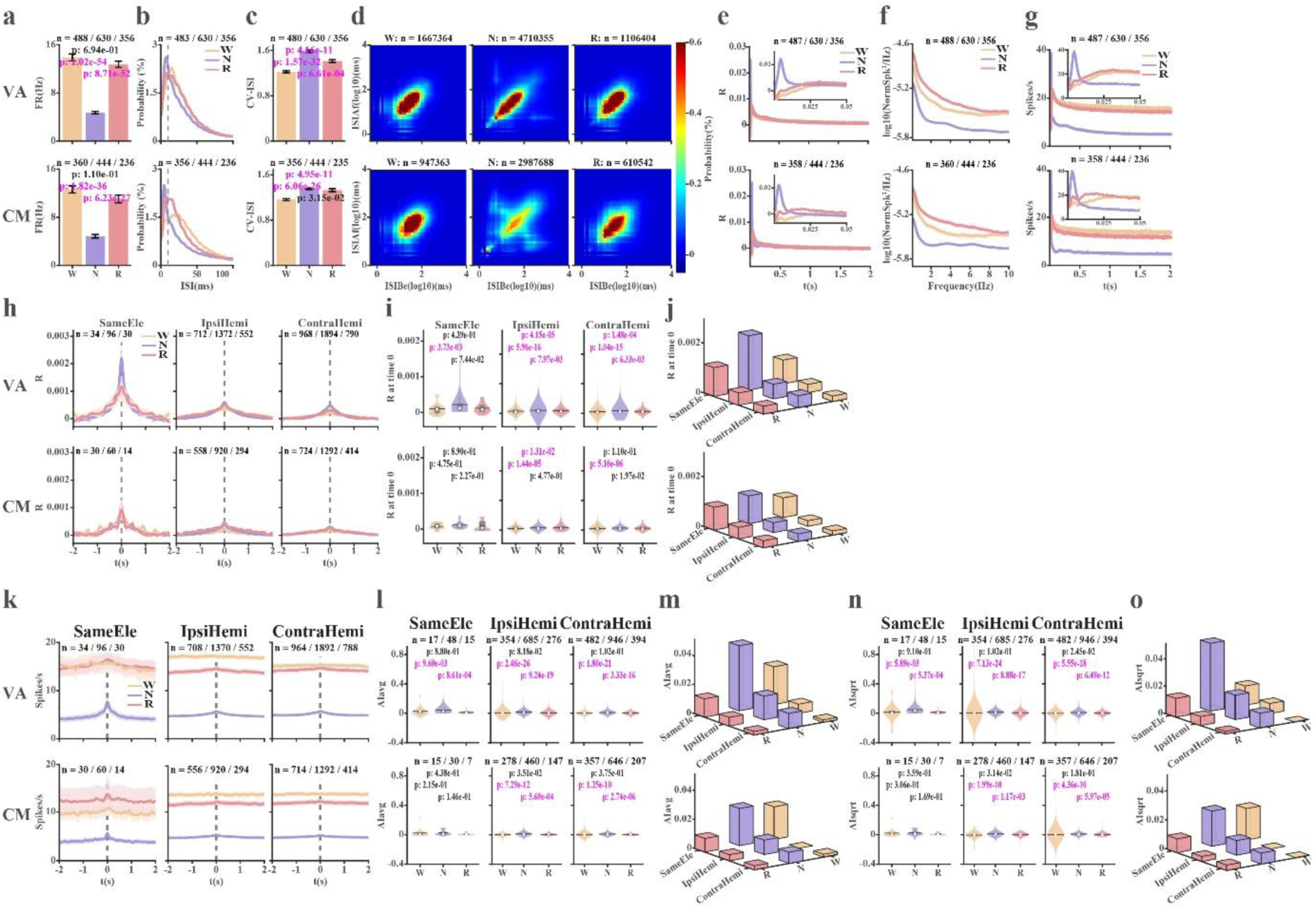
Analysis of thalamic spike trains after removing bursts (tonic activity). Binary (0/1) spike (isolation quality > 0.7) trains after removing detected bursts (by the maximum interval method). **a**, Firing rate (FR). **b**, inter spike interval (ISI). **c**, coefficient of variation of ISI (CV-ISI). **d**, ISI return map (ISIAf, ISI(n+1) vs ISIBe, ISI(n)). **e**, Average correlation coefficient (R) auto-correlation histograms for VA and CM tonic spike trains (lag = 2 s). The inset zooms in on the auto-correlation histograms near time zero (lag = 50 ms). **f**, Power spectrum density of tonic activity. **g**, Average conditional discharge rate (spikes/s) auto-correlation histograms for VA and CM tonic spike trains (lag = 2 s). Insets – autocorrelation histograms with a lag of 50 ms. **h**, Average correlation coefficient (R) cross-correlation histograms for thalamic neuron pairs recorded by the same electrode (SameEle), different electrodes in the ipsi- and contra-lateral hemisphere (IpsiHemi and ContraHemi). **i**, Violin plots illustrating the distribution of R values at time zero. The number of R values at time zero is the same as that of neuron pairs with repetition shown in **h**. **j**, 3D-bar plots showing the average R value at time zero as a function of vigilance state and distance. **k**, Average conditional discharge rate cross-correlation histograms of thalamic neuron pairs from the same electrode, ipsilateral hemisphere, or contralateral hemisphere. **l**, Violin plots illustrating the distribution of AI_avg_ values (equation 1). **m**, 3D-bar plots demonstrating the average AI_avg_ value as a function of vigilance state and distance. **n**, Violin plots showing the distribution of AI_sqrt_ values (equation 2). **o**, 3D-bar plots illustrating the average AI_sqrt_ values as a function of vigilance state and distance. Data is shown as mean ± SEM in subplots **e**-**h**, and **k**. The n indicates the number of neurons in **a-c** and **e-g**, ISI pairs in **d**, neuron pairs with repetition in **h** and **k**, and neuron pairs without repetition in **l** and **n**. A Bonferroni-corrected (p < 0.05/3) Wilcoxon rank sum test was used to calculate the statistical significance of different R values at time 0 (Subplot **i**) and AI (Subplots **l** and **n**). P-value indicating statistically significant difference is marked in magenta. Otherwise, it is marked in black. Abbreviations and color coding as in Fig. S1. Related to Figs 1, 2, and 3.

**Figure S7.**
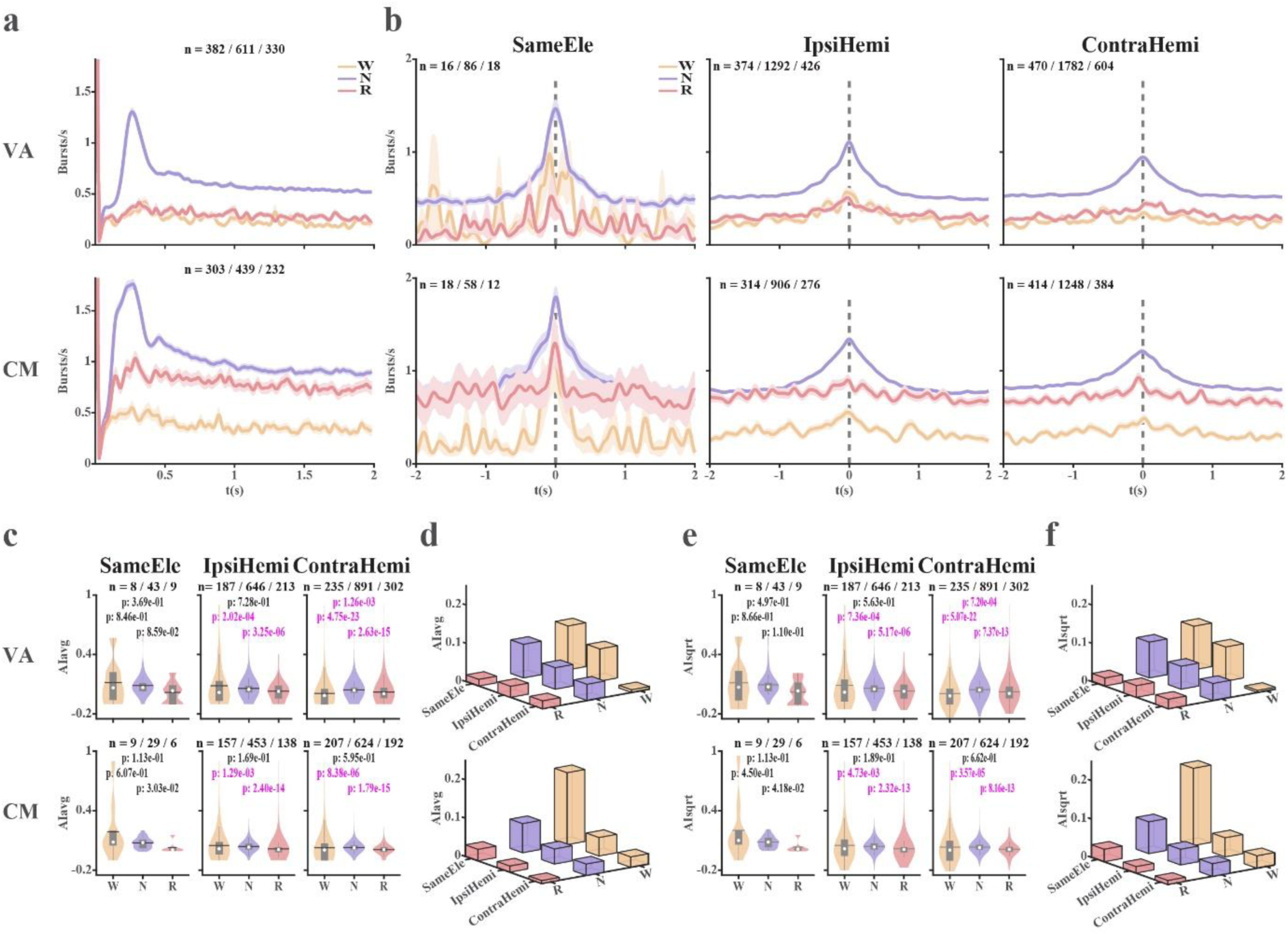
Auto- and cross-correlation conditional discharge rate histograms, and association indices of thalamic bursts. Bursts (detected by the maximum interval method) are treated as a single event at the time of their first spike. Auto and cross-correlation histograms of thalamic binary (0/1) burst trains are calculated as the conditional discharge rate following a burst at time zero. **a**, Conditional discharge rate auto-correlation histograms of burst trains. **b**, Conditional discharge rate cross-correlation histograms of burst trains. **c**, Association-index (AI, Fraction of bursts contributed by common-input mechanism out of total bursts) – average method (equation 1). Distributions of AI values are shown in violin plots. **d**, 3D-bar plots show average AI_avg_ values as a function of the vigilance stage and distance. **e**, AI of thalamic bursts – Sqrt method (equation 2). Same conventions as in subplot **c**. **f**, 3D-bar plots show average AI_sqrt_ values as a function of the vigilance stage and distance. Data is shown in mean ± SEM in **a** and **b**. The n indicates the number of neurons in **a**, neuron pairs with repetition in **b**, and neuron pairs without repetition in **c** and **e**. The Wilcoxon rank sum test was used to calculate the statistical significance of different AI values (Subplots **c** and **e**). The statistical significance of differences is corrected by the Bonferroni method, p < 0.05/3. The p-values indicating significant difference are marked in magenta. Otherwise, they are marked in black. Abbreviations and colour coding as in Fig. S1. Related to Figs. 2 and 3.

**Figure S8.**
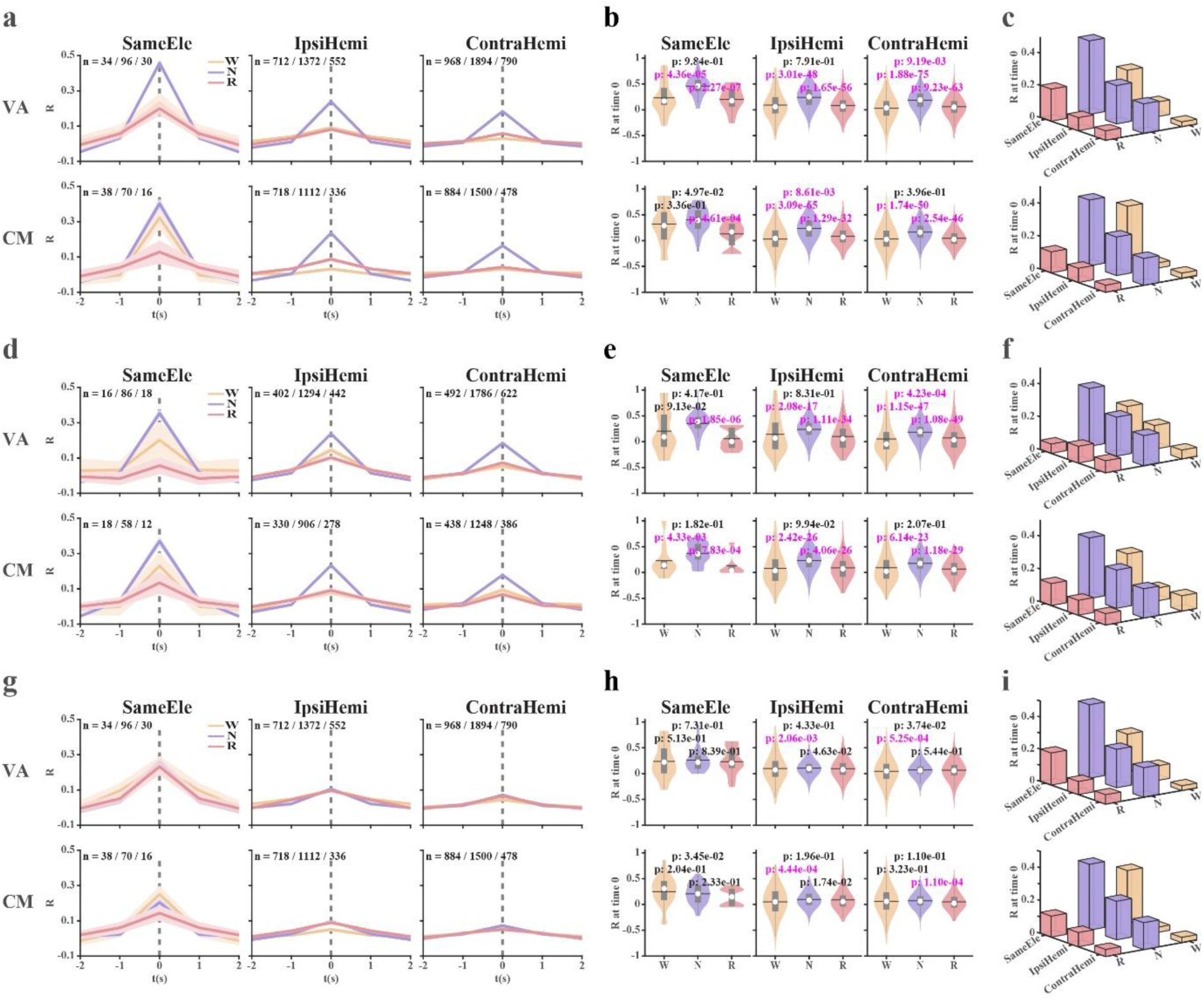
Cross-correlation histograms of spike trains, burst trains, and tonic activity (Time resolution = 1 s) across vigilance states. The isolation score of neural activity is larger than 0.7, and bursts are detected with the maximum interval (MI) method. **a-c**, All spike trains (single unit activity) are analysed. The average cross-correlation histograms of neuron pairs recorded from the same electrode, or different electrodes in the ipsilateral or contralateral hemisphere, are shown in **a**. Violin plots of the distribution of R values at time 0 gained from **a** are demonstrated in **b**. The number of time-zero R values is the same as that of neuron pairs with repetition shown in **a.** 3D-bar plots of the average R values at time zero as a function of vigilance states and distance (**c**). **d-f** and **g-i** show the same analysis for burst trains and for tonic activity (spike trains after removal of detected bursts), respectively. Data is shown in mean ± SEM, and n indicates the number of neuron pairs with repetition in **a**, **d**, and **g**. The statistically significant differences in R values at time zero among vigilance stages in **b**, **e**, and **h** are tested by Bonferroni-corrected Wilcoxon rank sum test (p<0.05/3). P values marked in magenta indicate significant differences; otherwise, they are black. Abbreviations and color coding as in Fig. S1. Related to Figs. 1**l-n**, 3**b-d** and S6**c-e**.

**Figure S9.**
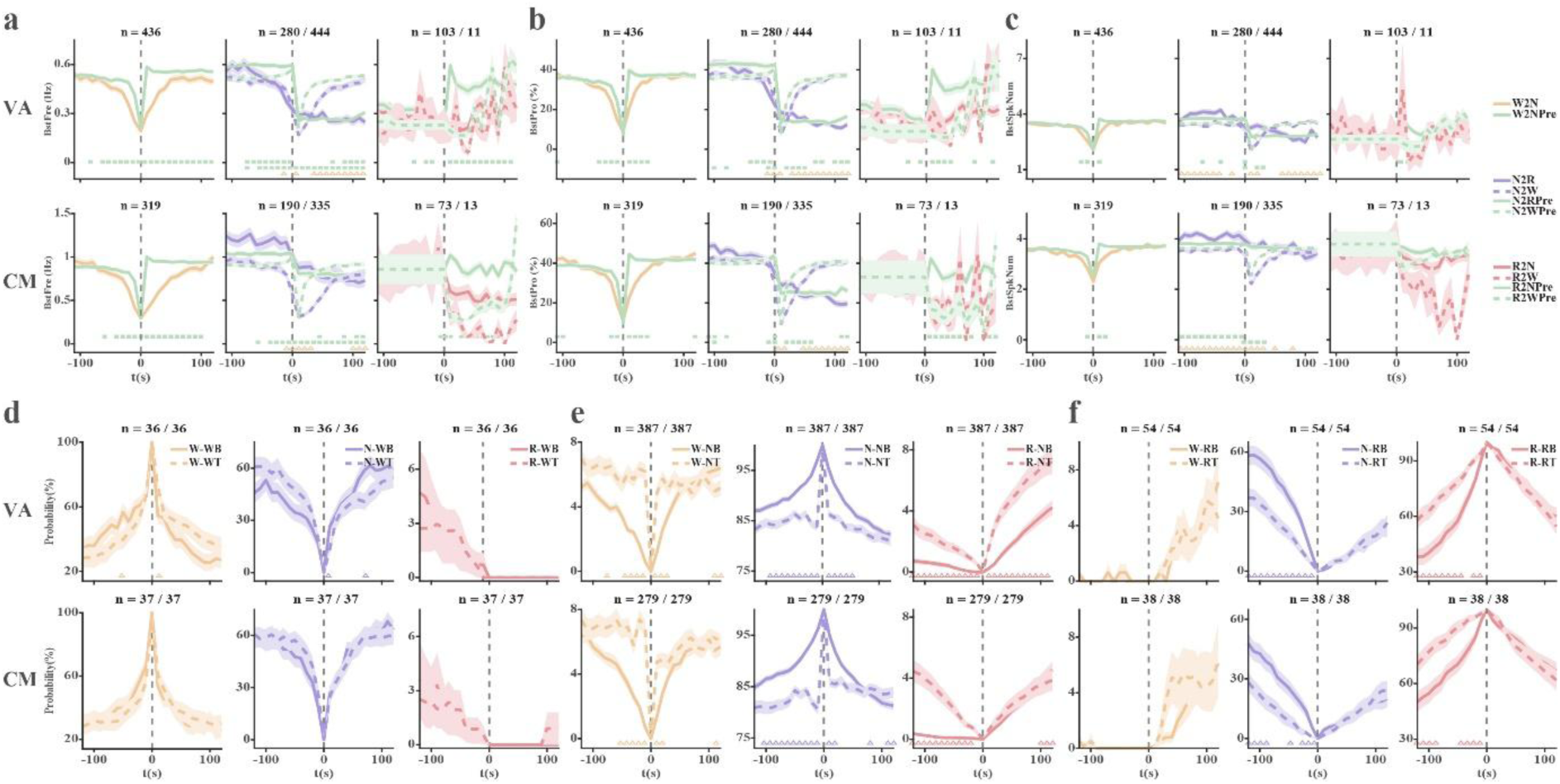
Basic analysis for vigilance stage transitions aligned with burst features and vigilance states aligned with firing mode (burst/tonic). **a-c**, The burst frequency (BstFre, **a**), fraction of spikes in bursts per 10 s total spikes (BstPro, **b**), and the average number of spikes per burst (BstSpkNum, **c**) aligned to different vigilance stages transition (e.g., W2N is the transition from wakefulness to NREM, and its corresponding predicted burst feature is W2NPre). There is no available data on the transition from wakefulness to REM sleep. The statistical difference between the actual and predicted values is tested using a Bonferroni-corrected Wilcoxon signed rank (p<0.05/24). Significant difference is marked by a green square (It indicates a significant difference between solid lines in the first row, and a significant difference between dashed lines in the second row). The statistical significance of differences between the transition to different vigilance stages from the same state is tested by Bonferroni corrected Wilcoxon rank sum (p<0.05/24), and their significant differences are marked by triangles coloured by corresponding sleep stages at time 0. **d-f**, The vigilance stages are aligned to segments with tonic or bursting segments combined with sleep states (Wakefulness in **d**, NREM in **e** and REM in **f**), e.g., NREM-burst (NB), NREM-Tonic (NT); W-NB indicates the percentage of wake aligned with NREM-Burst combination segment. The Bonferroni-corrected (p < 0.05/25) Wilcoxon signed rank is used to calculate the statistical significance of the difference. The values at time 0 are not compared. The significant difference is marked in color-coded triangles. Data is shown as mean ± SEM. The n indicates the number of neurons. The number of the predicted burst features is the same as that of their corresponding actual burst features. Abbreviations and color coding as in Fig. S1. Related to Fig. 4**a**, **b**.

